# NINJ1 mediates necrosis of *Mycobacterium tuberculosis* infected human macrophages

**DOI:** 10.1101/2025.11.04.685750

**Authors:** Ragnhild SR Sætra, Mathilde Hansen, Shirin Kappelhoff, Marit Bugge, Anne Marstad, Liv Ryan, Pascal Devant, Jonathan C Kagan, Kristine Freude, Kai S Beckwith, Trude Helen Flo

## Abstract

Lytic cell death contributes to tissue damage and dissemination during *Mycobacterium tuberculosis* (Mtb) infection, yet the mechanisms governing plasma membrane rupture (PMR) remain incompletely defined. Here, we identify Ninjurin-1 (NINJ1), a mediator of PMR, as a central effector of macrophage lysis during Mtb infection. Mtb infection induced NINJ1 oligomerization, and genetic deletion or pharmacological inhibition of NINJ1 markedly reduced lactate dehydrogenase (LDH) release. Cytokine secretion was largely preserved in NINJ1-deficient macrophages, with the exception of CXCL10. Unexpectedly, inhibition of pyroptosis, apoptosis, necroptosis, and ferroptosis - individually or in combination - did not prevent NINJ1 activation or PMR, indicating that Mtb-induced membrane rupture proceeds independently of canonical regulated cell death pathways. However, NINJ1 oligomerization and PMR required the Mtb ESX-1 secretion system, identifying a bacterial virulence determinant as a critical upstream trigger. Although osmoprotective PEG partially reduced LDH release, neither calcium signaling nor cell swelling accounted for NINJ1-dependent PMR in Mtb-infected macrophages. Together, these findings establish NINJ1 as a key executioner of Mtb-induced lytic cell death and reveal an ESX-1-dependent pathway of PMR that is uncoupled from canonical host cell death programs.

**Teaser:** NINJ1 mediates plasma membrane rupture of macrophages infected with *Mycobacterium tuberculosis*

## INTRODUCTION

Macrophages are the first cells infected by *Mycobacterium tuberculosis* (Mtb) when it enters the human lung and provide an important niche for the bacterium over the course of infection(1). After uptake, Mtb modulates intracellular trafficking to survive and replicate in phagosomes and occasionally translocate into the cytosol(2–4). Phagosomal rupture and the presence of Mtb in the cytosol is sensed by pattern recognition receptors and triggers signaling cascades culminating in the release of inflammatory cytokines and interferons, as well as inflammatory cell death(1, 2, 5). The induction of cell death is linked to virulence properties of the Mtb cell wall, and especially the capacity to damage host phagosome- and plasma membranes (PM) through activities of the type VII secretion system, ESX-1, and the cell wall lipid phthiocerol dimycocerosate (PDIM)(3, 4, 6–12).

Mtb-infection drives multiple modes of regulated cell death (RCD)(13, 14) including apoptosis(15, 16), pyroptosis(9, 17–19), necroptosis(20), and ferroptosis(21–24). Apoptosis involves activation of caspases 8/9/7/3 by death receptors or intracellular stress, leading to cellular degradation and formation of apoptotic bodies cleared by phagocytes (efferocytosis). Apoptosis is generally non-inflammatory unless efferocytosis is insufficient and the cells progress to secondary necrosis (post-apoptotic lysis)(25). Pyroptosis, necroptosis and ferroptosis provoke immune responses due to their lytic nature, releasing cytosolic content including microbes and damage-associated molecular patterns (DAMPs), into the tissue(26). Necroptosis is triggered by death receptors or Toll-like receptors (TLRs) in cases where the apoptotic caspase 8 is inhibited, whereby Receptor-interacting protein kinases (RIPK)1/3 form a complex that phosphorylates Mixed lineage kinase domain like pseudokinase (MLKL) which oligomerizes and forms plasma membrane lesions(27). Many pathways activate pyroptosis, but canonically it involves the assembly of cytosolic inflammasomes and activation of caspases 1/4/5, and results in the formation of gasdermin pores in the PM for release of interleukin-(IL)-1β and IL-18(28). Ferroptosis is caspase independent and results from accumulation of lipid peroxides in the PM in situations of iron overload, inhibition of glutathione peroxidase 4 (GPX-4), or depletion of cellular antioxidants(29).

In tuberculosis, apoptosis is generally considered favorable by eliminating the replicative niche of Mtb and causing little tissue damage, while necrotic cell death may benefit the bacterium by promoting spread(12, 13, 16, 30) (14). However, concomitant activation of pattern recognition receptors in dying cells contributes to the resulting immune response and differ depending on the cell type and cell death modality engaged. For instance, pyroptosis is highly inflammatory due to the release of IL-1β and IL-18 - key cytokines involved in protective immunity against Mtb(31). On the other hand, Mtb delivers several bacterial effectors into host cells that interfere with cell death pathways directly, including pyroptosis (PknF(32), PtpB(33)), apoptosis(34) and ferroptosis (PtpA(24)), or indirectly by interfering with membrane repair(35). The circumstances under which RCD pathways are activated in tuberculosis, as well as their significance, is not understood.

Terminal lysis in necrotic cell death was long considered a passive consequence of pathway-specific membrane permeabilization e.g., by gasdermins(36), MLKL(27), or lipid peroxidation(37), leading to osmotic swelling and plasma membrane rupture (PMR). Recently, the small membrane protein ninjurin 1 (NINJ1) was found to actively mediate PMR and cell lysis in several cell death modalities including pyroptosis, apoptosis, and from bacterial pore-forming toxins(38). Importantly, IL-1β secretion was intact in NINJ1-deficient pyroptotic cells, placing NINJ1 downstream of gasdermin D (GSDMD) pore formation, while NINJ1-mediated lysis was required for release of larger danger molecules like LDH and HMGB1(38). Cell lysis during necroptosis was less dependent on NINJ1(38) and recently suggested to be mediated by SIGLEC12(39). Later studies of ferroptosis have suggested a partial or full dependence of NINJ1 for PMR(37, 40, 41). NINJ1 is proposed to exist as monomers and autoinhibited dimers in the PM of resting cells(42). Upon activation, the dimers dissociate and NINJ1 monomers oligomerize to form large, filamentous structures that eventually mediate PMR(38, 43–47). Structural studies of NINJ1 have proposed distinct models for how PMR is executed: via pore formation or assembly and release of NINJ1-encircled membrane disks(43, 46), or filamentous zipper-like assemblies that open upon cell swelling due to increased membrane tension or mechanical strain(40, 44, 45, 47, 48). However, the mechanisms controlling NINJ1 activation are not known. Recent data, some still in open archives, suggest that changes in the biophysical properties of the PM or Ca^2+^-dependent phospholipid scrambling can trigger NINJ1 oligomerization(47, 49, 50).

Only a few studies have explored the role of NINJ1 in infection: in mice, NINJ1 deficiency increased the susceptibility to infection with *Clostridium*(38) or *Yersinia*(51), and NINJ1 has been found to play a role in *Salmonella* infection of mouse(38) or human(52) macrophages. Here, we show that NINJ1 mediates PMR in Mtb-infected macrophages. NINJ1 activation required the Mtb ESX-1 secretion system but occurred independently of the RCD pathways apoptosis, pyroptosis, necroptosis, and ferroptosis, suggesting that pathogen-driven membrane perturbations act as the upstream trigger. Given the multiple cell death pathways engaged by Mtb, targeting a common mediator of necrosis such as NINJ1 may offer a promising strategy to limit inflammation, tissue damage, and Mtb dissemination.

## RESULTS

### Infection with *Mycobacterium tuberculosis* activates NINJ1 oligomerization and NINJ1-dependent plasma membrane rupture of human macrophages

To investigate the role of NINJ1 in Mtb-induced lytic cell death, we infected human iPSC-derived macrophages (iPSDMs) with the BSL2-adapted auxotrophic Mtb strain H37Rv mc^2^6206::BFP (hereafter referred to as Mtb mc^2^6206)(53), which retains key virulence traits including ESX-1(9). CRISPR/Cas9-mediated knockout of NINJ1 significantly reduced LDH-release from infected iPSDMs (Fig. 1A and Fig. S1A). Accordingly, inhibition of NINJ1 oligomerization using glycine (54) reduced LDH release in Mtb-infected WT but not NINJ1 KO cells (Fig. 1A). Native-PAGE revealed a shift of NINJ1 monomers/dimers to higher-order oligomers upon infection, confirming that NINJ1 oligomerization was triggered by Mtb-infection (Fig. 1B). Similar results were obtained when NINJ1 oligomerization dynamics at the PM was monitored via live-cell total internal reflection fluorescence (TIRF) microscopy of NINJ1 KO THP-1 macrophages reconstituted with hNINJ1-mNeonGreen (THP-1 NINJ1-mNG cells) (Fig. 1C-F, Movie S1). At resting conditions, NINJ1-mNG was homogeneously distributed across the PM. Upon Mtb infection, NINJ1 clustered into bright, distinct puncta over 3–20 h. This process appeared slower than NINJ1 oligomerization in iPSDMs (4 h) but was prevented by glycine pretreatment in both cell types. Once initiated, however, NINJ1 oligomerization proceeded rapidly, with a half-time shorter than the 5 min imaging/frame interval (Fig. 1F). NINJ1 oligomerization was accompanied by nuclear accumulation of the membrane-impermeant dye DRAQ7 within the 5 min. frame interval (Fig. 1E, F), indicative of plasma membrane permeabilization. Collectively, these findings demonstrate that NINJ1 oligomerizes during Mtb-infection and mediates PMR in infected macrophages.

**Figure 1:**
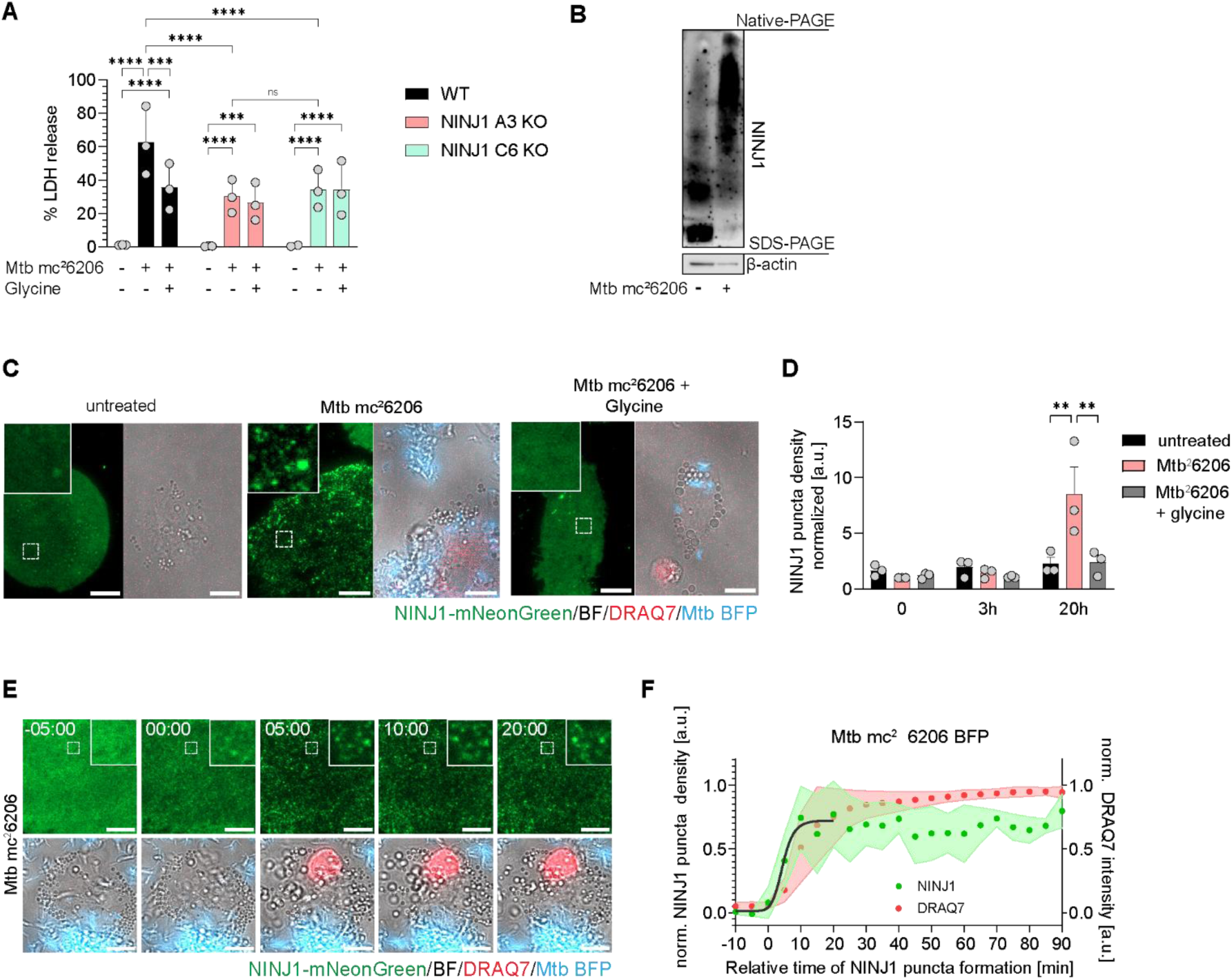
NINJ1 oligomerizes and mediates PMR in Mtb-infected human macrophages. (**A, B**) LDH release (A) and NINJ1 oligomerization (Native-PAGE) (B) from WT and NINJ1 KO iPSDMs pretreated with glycine and infected with Mtb mc²6206, MOI 20 for 4h. Data are means (bars) ± s.e.m from N = 3 independent experiments (circles), each with three replicates per condition (A). Significant differences are indicated as *P<0.05; **P<0.01; ****P<0.0001 by RM two-way Anova with Šídák’s multiple comparisons test. (**C**) Representative images of NINJ1 KO THP-1 cells expressing hNINJ1-mNG (THP1-NINJ1-mNeonGreen) with or without glycine pre-treatment and infection with Mtb mc^2^6206-BFP for 20h. TIRF (mNG), widefield (DRAQ7 and BFP) and brightfield microscopy images of NINJ1-mNG (green), DRAQ7 (red) and Mtb (blue), and morphology, respectively. Scale bars 10 µm. (**D**) Quantification of NINJ1-mNG puncta densities at 0h, 3h and 20h of infection (Mtb) as represented in C. Data are means (bars) ± s.e.m from N = 3 independent experiments (circles), each including analysis of 4-15 cells per condition. Significant differences are indicated as *P<0.05; **P<0.01; ****P<0.0001 by RM two-way Anova with Tukey’s multiple comparisons test. (**E**) Representative time-lapse images of NINJ1 clustering (TIRF, green) and DRAQ7 uptake (red) upon Mtb mc^2^6206-BFP (blue) infection in a THP1-NINJ1-mNeonGreen cell. Scale bars 10 µm. (**F**) Kinetics of the increase of NINJ1 puncta densities and DRAQ7 intensities upon Mtb mc^2^6206-BFP infection monitored with a frame rate of 5 min and temporally aligned by start of NINJ1 clustering (t = 0). Density values were normalized to the highest value for each cell and aligned by the start of NINJ1 oligomerization. Data are means ± s.d. with sigmoidal fits for NINJ1-mNG puncta density increase (green). N = 4 experiments, 10 cells. Time in min.

### NINJ1 oligomerizes in apoptosis, pyroptosis, necroptosis and ferroptosis, but is only required for LDH-release during pyroptosis and post-apoptotic lysis

Multiple cell death modalities occur in Mtb-infected macrophages(13, 55). To investigate the contribution of individual RCD modalities to NINJ1-mediated lysis of Mtb-infected macrophages, we compared NINJ1 activation and -dependency in apoptosis, pyroptosis, necroptosis and ferroptosis. First, the phenotype and kinetics of NINJ1 oligomerization was monitored in THP-1 NINJ1-mNG cells and iBMDM hNINJ1-mNG cells upon treatment with different RCD stimuli: Venetoclax (apoptosis), LPS+Nigericin (pyroptosis), and RSL-3 (ferroptosis). Induction of all three RCD pathways triggered NINJ1 oligomerization into distinct, bright puncta at the PM within 4 h and with similar densities (Fig. 2A, B and Fig. S2A-C, E, F). Glycine pretreatment abolished NINJ1 oligomerization in all conditions (Fig. 2A, B, Fig. S2A-C, G) like in Mtb-infected cells (Fig. 1). Oligomerization proceeded rapidly upon induction of all pathways in both cell types, with half-times (t_50_) ranging from 0.5 to 1.17 min (apoptosis: 0.5 ± 0.12 min, ferroptosis: 0.89 ± 0.06 min, and pyroptosis: 1.17 ± 0.12 min) (Fig. 2C and Fig. S2D-F and Movie S2-4). NINJ1 oligomerization was accompanied by influx of DRAQ7 (Fig. S2D-F). These data suggest that the NINJ1 puncta phenotype is largely independent of the cell death trigger and therefore insufficient to distinguish between RCD modalities potentially contributing to Mtb-induced NINJ1 activation.

**Figure 2:**
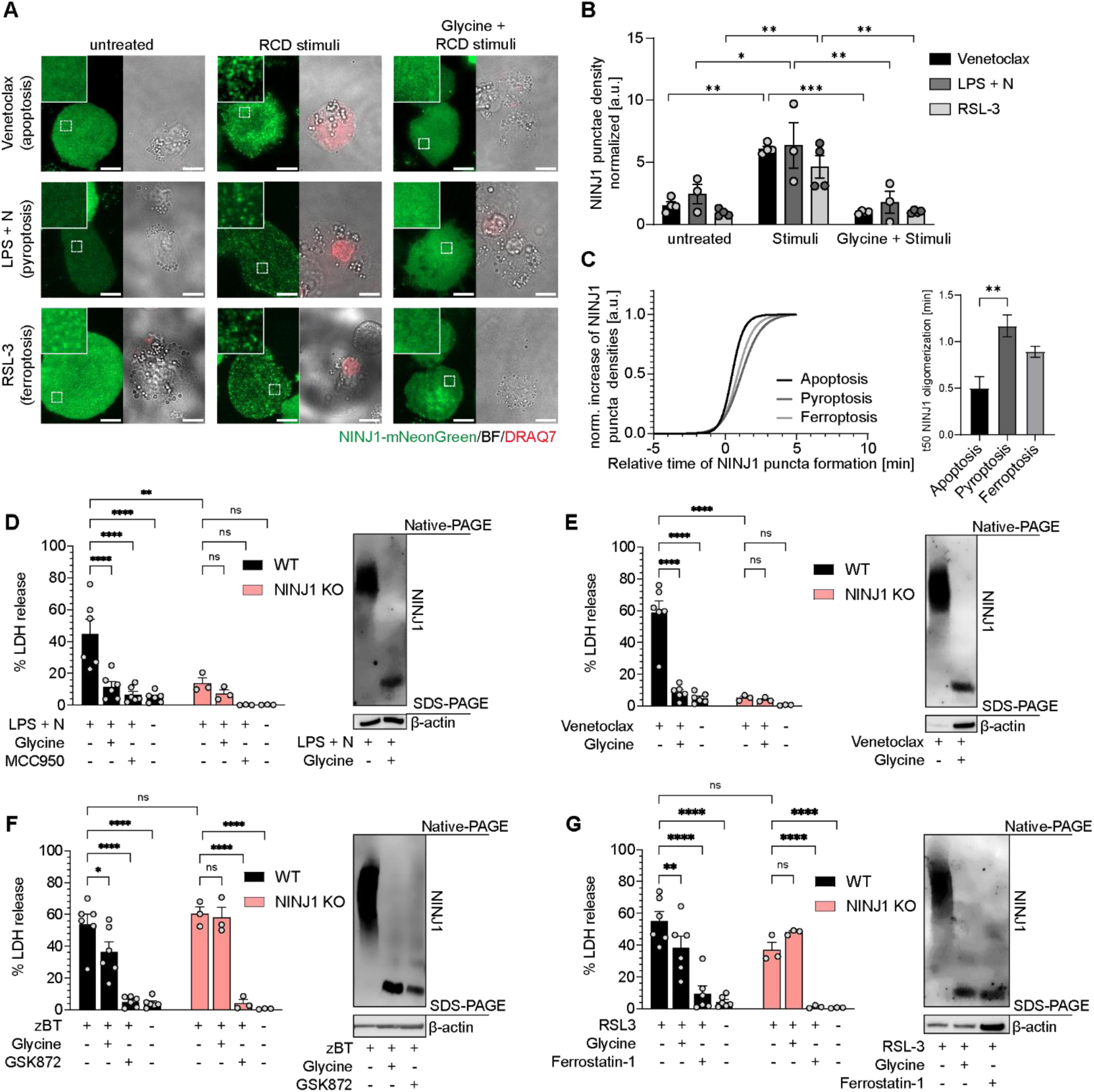
NINJ1 oligomerization is induced during pyroptosis, apoptosis, necroptosis and ferroptosis but is only required for plasma membrane rupture of pyroptotic and apoptotic human macrophages. (**A**) Representative images of THP1-NINJ1-mNeonGreen cells pre-treated with or without glycine and treated with cell death stimuli venetoclax (apoptosis), LPS + nigericin (LPS+N, pyroptosis) or RSL-3 (ferroptosis) for 4h. TIRF (mNG, green), widefield (DRAQ7, red) and brightfield microscopy images. Scale bars 10 µm. (**B**) Quantification of NINJ1-mNG puncta densities at 4h (Venetoclax, LPS+N, RSL-3) as represented in (A). Data are means (bars) ± s.e.m from N = 3-4 independent experiments (circles), each including analysis of 4-15 cells per condition. Significant differences are indicated as *P<0,05 by RM two-way Anova with Tukey’s multiple comparisons test. (**C**) Normalized sigmoidal fits of NINJ1 puncta density increase (left) and t_50_ values (right, x value where y=50% max.) of apoptosis (Venetoclax), pyroptosis (LPS+N) or ferroptosis (RSL-3) induction in THP1-NINJ1-mNeonGreen cells monitored with frame rates of 8-10 sec. N = 3 experiments, 8-12 cells. Individual kinetic curves and representative images for RCD induction are shown in Supplementary Fig. 2. (**D**-**G**) NINJ1 oligomerization (Native-PAGE) and LDH release from WT and NINJ1 KO iPSDMs pretreated with or without glycine or inhibitors as indicated and cell death stimuli for 4h: (D) pyroptosis – LPS and nigericin (LPS+N) w/wo NLRP3 inhibitor MCC950; (E) apoptosis – venetoclax; (F) necroptosis – Z-VAD-FMK+BV-6+TNF-α (zBT) w/wo RIPK3 inhibitor GSK’872; (G) ferroptosis – RSL-3 w/wo Ferrostatin-1. Data are means (bars) ± s.e.m. of at least N = 3 independent experiments (circles), each with three replicates per condition. *P<0.05; **P<0.01; ****P<0.0001 by RM two-way Anova with Tukey’s multiple comparisons test of treatments within each cell line and stimuli between cell lines.

We next examined the NINJ1-dependency of PMR in apoptosis, pyroptosis, necroptosis, and ferroptosis by treating WT and NINJ1 KO iPSDMs with agonists for each modality with or without specific inhibitors MCC950 (NLRP3, pyroptosis), GSK’872 (RIPK3, necroptosis), Ferrostatin-1 (radical-trapping antioxidant, ferroptosis) or glycine. Like in THP-1s and iBMDMs, NINJ1 oligomerized upon ligand-induced apoptosis, pyroptosis, necroptosis and ferroptosis in iPSDMs (Fig. 2D-G). Consistent with our earlier work(54), glycine or NINJ1 deletion inhibited LDH-release during pyroptosis comparable to direct pathway inhibition by MCC950 (Fig. 2D and Fig. S2H, S3A). Similarly, glycine as well as NINJ1 deletion abolished LDH release during apoptosis/post-apoptotic lysis (Fig. 2E and Fig. S3B). GSDMD KO iPSDMs were generated (Fig. S3E-H) and as expected, LDH-release was almost abolished during pyroptosis but not apoptosis (Fig. S3E, F), indicating PMR is GSDMD-dependent in pyroptosis and GSDMD-independent in post-apoptotic lysis. Glycine had no added effect in NINJ1 KO iPSDMs (Fig. 2D, E and Fig. S3A, B), confirming that PMR in pyroptosis and apoptosis is NINJ1-dependent. Secretion of IL-1β and IL-18 from pyroptotic cells was inhibited by MCC950, but not glycine, while TNF-α and IL-6 secretion remained unaffected (Fig. S4A). Apoptosis induced minimal cytokine production (Fig. S4B). Strikingly, LDH release in necroptotic and ferroptotic iPSDMs was abolished by pathway-specific inhibitors (GSK’872 and Ferrostatin-1, respectively), but glycine only modestly reduced LDH release, and NINJ1 deletion had no effect (Fig. 2F, G and Fig. S3C, D). These findings suggest that NINJ1 is activated, but not required, for PMR during necroptosis and ferroptosis. They further indicate that neither necroptosis nor ferroptosis are main contributors to PMR in Mtb-infected macrophages. Cytokine secretion was low in necroptotic and ferroptotic iPSDMs (Fig. S4C, D). Collectively, these data suggest that while NINJ1 oligomerizes during all RCD modalities, it is needed for PMR only during pyroptosis and apoptosis and not during necroptosis or ferroptosis.

### Mtb-induced NINJ1 oligomerization and PMR is independent of canonical host cell death programs apoptosis, pyroptosis, necroptosis or ferroptosis

RCD pathway-specific inhibitors were next used to assess whether any individual pathway contributes to NINJ1-mediated PMR and inflammatory cytokine release during Mtb-infection. iPSDMs were pretreated with glycine, MCC950, Z-VAD-FMK (zVAD, pan-caspase inhibitor), GSK’872 or Ferrostatin-1 before infection with Mtb mc^2^6206 or H37Rv. LDH-release was unaffected by individual inhibitors (Fig. 3A and Fig. S5A), despite their efficacy in blocking ligand-induced NINJ1 oligomerization and LDH-release (Fig. 2). Only glycine reduced LDH-release upon Mtb-infection (Fig. 3A and Fig. S5A). We also tested 2-Bromohexadecanoic acid (2-BP), a general palmitoylation inhibitor, since palmitoylation is shown to modulate different RCD pathways(56–62). However, 2-BP had no effect on LDH-release in WT or NINJ1 KO iPSDMs (Fig. S5B). While LDH-release remained unaffected by RCD inhibitors during Mtb infection, cytokine release was modulated. IL-1β release was strongly inhibited by zVAD and MCC950 during Mtb mc^2^6206 and H37Rv infection (Fig. 3B and Fig. S5C, D), confirming our previous finding that the NLRP3-caspase-1-GSDMD axis is the main mechanism of IL-1β release from Mtb-infected human macrophages(9). zVAD and GSK’872 both reduced the secretion of IL-6, IL-10, IL-1Ra and IFN-β from Mtb mc^2^6206-infected iPSDMs (Fig. S6), indicating signaling via caspases and RIPK3.

**Figure 3:**
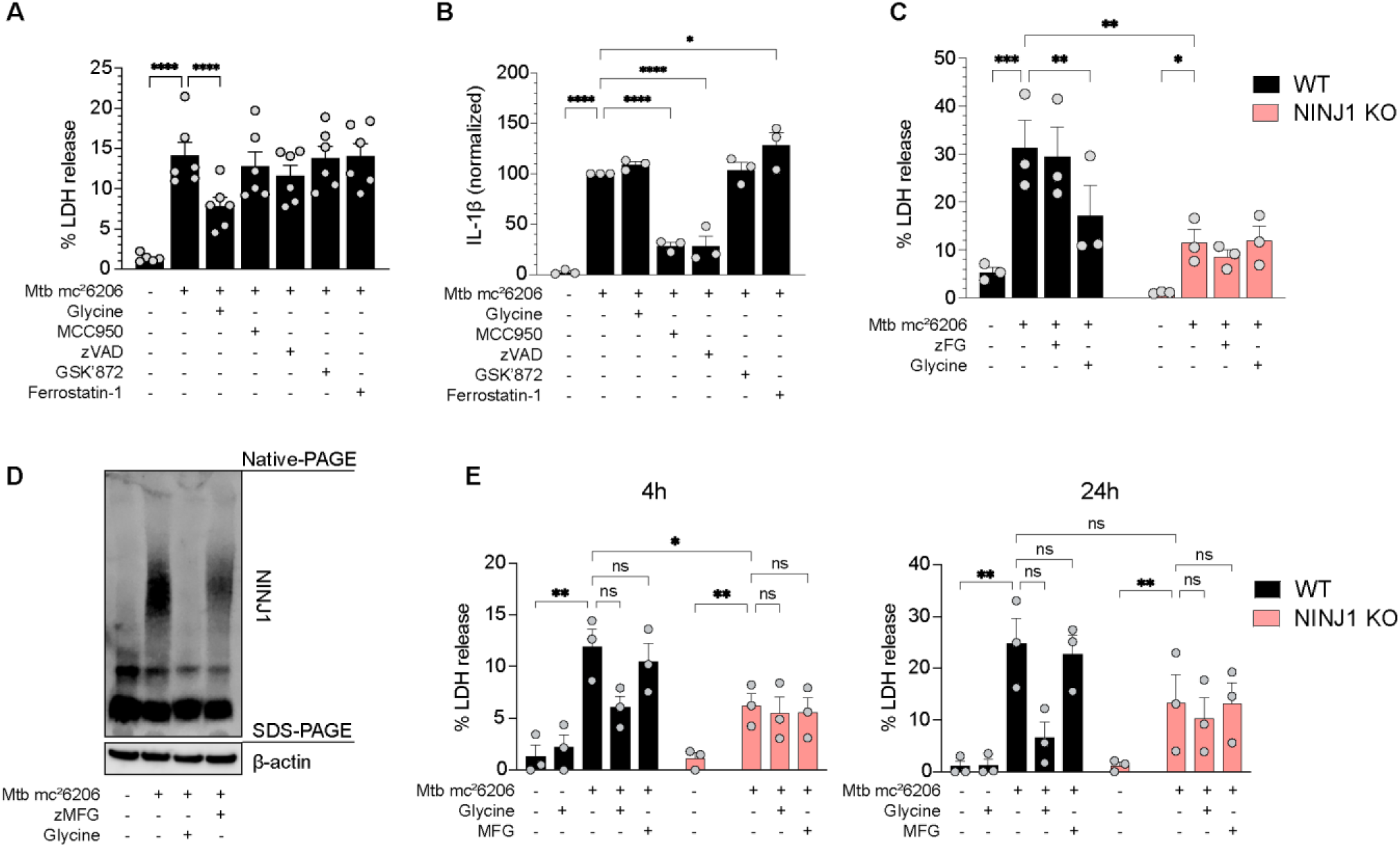
NINJ1 oligomerization and plasma membrane rupture in Mtb-infected macrophages are not affected by cell death specific inhibitors. (**A, B**) LDH (A) and IL-1β release (B) from iPSDM WTs pretreated with glycine and cell death inhibitors (MCC950, Z-VAD-FMK, GSK’872 and Ferrostatin-1) and infected with Mtb mc²6206, MOI 20 for 4h. Data are means (bars) ± s.e.m. (A) or means ± s.e.m. normalized against the Mtb mc^2^6206 infected sample (B) of N = 6 (A) or N = 3 (B) independent experiments (circles), each with three replicates per condition. *P<0.05; **P<0.01; ***P<0.0001 by RM one-way ANOVA with Dunnett’s multiple comparisons test against the infected sample. (**C**) LDH release from WT and NINJ1 KO iPSDMs pretreated with or without glycine or a cocktail of inhibitors (zFG) including Z-VAD-FMK, Ferrostatin-1 and GSK’872 and infected with Mtb mc²6206, MOI 20 for 4h. Data are means (bars) ± s.e.m. of N = 3 independent experiments (circles), each with three replicates per condition. *P<0.05; **P<0.01; ****P<0.0001 by RM two-way ANOVA with Šídák’s multiple comparisons test of treatments within each cell line and stimuli between cell lines. (**D**) NINJ1 oligomerization (Native-PAGE) in WT iPSDMs pretreated with or without glycine or a cocktail of inhibitors (zMFG) including Z-VAD-FMK, MCC950, Ferrostatin-1 and GSK’872 and infected with Mtb mc²6206, MOI 20 for 4h. (**E**) LDH release from WT and NINJ1 KO iPSDMs pretreated with or without glycine or a cocktail of inhibitors (MFG) including MCC950, Ferrostatin-1 and GSK’872 before infection with Mtb mc²6206, MOI 2 for 4h or 24h. Data are means (bars) ± s.e.m. of N = 3 independent experiments (circles), each with three replicates per condition.. *P<0.05; **P<0.01; ****P<0.0001 by RM two-way ANOVA with Šídák’s multiple comparisons test of treatments within each cell line and stimuli between cell lines.

To test whether RCD pathway redundancy explains the failure of individual inhibitors to block LDH-release during Mtb infection(26), iPSDMs were pretreated with a cocktail of zVAD, Ferrostatin-1, and GSK’872 (zFG) before infection with Mtb mc²6206. Surprisingly, LDH-release remained unaffected by the zFG inhibitor cocktail and was only reduced by glycine or in NINJ1 KO iPSDMs (Fig. 3C). Similar trends were observed during Mtb H37Rv infection (Fig. S5E). Furthermore, Native-PAGE confirmed that NINJ1 oligomerization was prevented by glycine but not by zMFG (zFG + MCC950) (Fig. 3D), indicating that NINJ1 activation and PMR occurs independently of caspases, NLRP3, RIPK3 or lipid peroxidation during Mtb infection.

While zFG did not affect LDH-release, it significantly reduced the secretion of IL-1α, IL-1β, TNF-α, and IFN-α during Mtb mc²6206 infection (4h) in both WT and NINJ1 KO iPSDMs, with IL-6 and IL-10 showing a similar trend (Fig. S7A). IL-8 release was unaffected. Similar patterns were seen with Mtb H37Rv infection (24h), though not statistically significant (Fig. S7B). Mtb-induced cytokine secretion was largely unaffected by NINJ1 inhibition or deletion, except for CXCL10, which was significantly reduced in NINJ1 KO iPSDMs during mc²6206 infection, with a similar trend for H37Rv or following glycine treatment in WT cells (Fig. S7A and B).

The preceding experiments were conducted at high multiplicity of infection (MOI 20), which favors rapid membrane damage and cell death(9, 12, 30, 63–65). We next tested whether lower MOI, where cytotoxicity is expected to develop more gradually, also induces NINJ1-dependent PMR and, if so, whether canonical RCD pathways are drivers. WT and NINJ1 KO iPSDMs were infected with Mtb mc²6206 at MOI 2 in the presence or absence of glycine or MFG (MCC950, Ferrostatin-1, and GSK’872). Low-MOI infection induced detectable LDH release after 4h (∼12%), which increased approximately twofold by 24h (Fig. 3E). LDH release was consistently reduced by ∼50% in NINJ1 KO iPSDMs and with glycine treatment, although statistical significance was reached only for NINJ1 deficiency at 4h. In contrast, pretreatment with the MFG inhibitor cocktail had no effect on LDH release in either WT or NINJ1 KO iPSDMs at either time point. Collectively, these findings show that NINJ1-dependent PMR occurs under both high- and low-burden Mtb infection in human macrophages and is independent of canonical apoptosis, pyroptosis, necroptosis, and ferroptosis pathways.

### NINJ1 activation is triggered by ESX-1-induced plasma membrane perturbations during Mtb infection

Mtb can cause ESX-1-dependent phagosomal membrane damage and direct, uptake-independent damage to the PM, leading to NLRP3 inflammasome activation and pyroptosis(9, 12). However, since Mtb-induced NINJ1 activation is independent of pyroptosis and other major RCD pathways, we tested whether extracellular Mtb can activate NINJ1 and if ESX-1 is required. Blocking phagocytosis with Cytochalasin D in iBMDM NINJ1-mNG cells prevented uptake of Mtb mc²6206 but did not inhibit NINJ1 oligomerization or DRAQ7 uptake (Fig. 4A-D and Movie S5 and S6). Cytochalasin D alone did not affect NINJ1 oligomerization or cell permeabilization (Fig.4A-C, E, F). In iPSDMs, inhibition of phagocytosis with Cytochalasin D slightly reduced LDH-release upon Mtb infection, but Native-PAGE revealed that NINJ1 oligomerization was unaffected (Fig. 4E, F). To assess the role of ESX-1, we infected cells with the RD-1-deficient mutant Mtb mc²6230, which lacks most of the ESX-1 secretion system(66). Unlike Mtb mc²6206, the RD-1 mutant mc²6230 strain failed to induce NINJ1 oligomerization and cell permeabilization in iBMDMs over 20h infection (Fig. 4A-D) and did not trigger NINJ1 oligomerization or LDH-release in iPSDMs (Fig. 4E, F). Together, these results show that Mtb-induced NINJ1 oligomerization and PMR is ESX-1 dependent and does not require uptake, suggesting that ESX-1-inflicted PM perturbations trigger NINJ1 activation.

**Figure 4:**
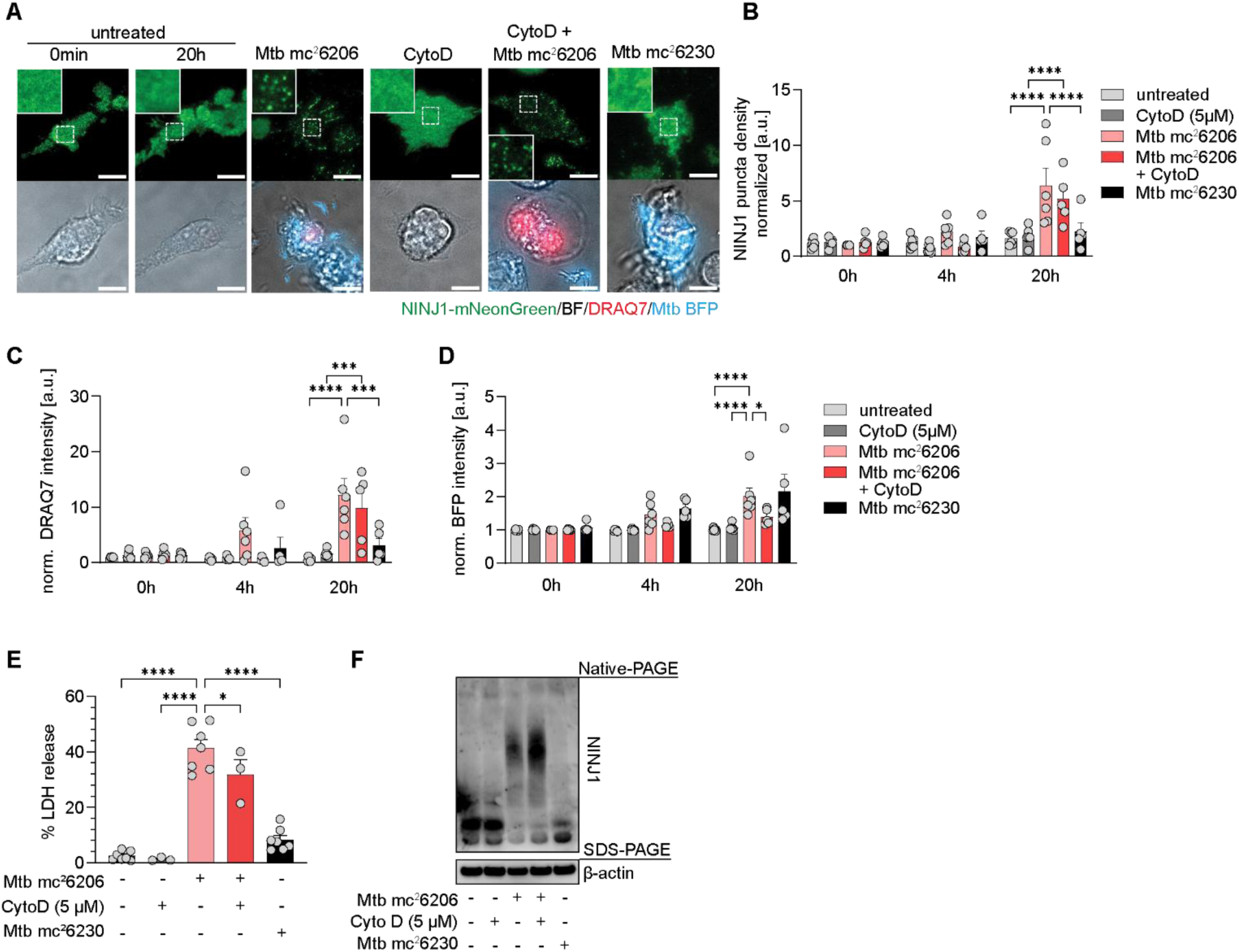
NINJ1 oligomerization and PMR is driven by ESX-1-induced plasma membrane damage. (**A-D**) NINJ1-mNG oligomerization in immortalized mouse NINJ1-KO BMDMs expressing fluorescently tagged human NINJ1 with or without 5 µM Cytochalasin D pretreatment to prevent phagocytosis followed by infection with Mtb mc²6206-BFP or the RD-1 deficient mutant Mtb mc^2^6230-BFP. Representative TIRF microscopy images (A) of hNINJ1-mNG (green), DRAQ7 uptake (red, WF) and bacteria (blue, WF) after 20h of infection and quantification of NINJ1 puncta densities (B), intracellular DRAQ7 (C) and BFP (D) after 0, 4 and 20h. Data are means (bars) ± s.e.m. from N = 5-6 independent experiments with 4-10 cells per condition each. Significant differences are indicated as *P<0.05; **P<0.01; ****P<0.0001 by RM two-way Anova with Šídák’s multiple comparisons test. Only significant differences within each timepoint are shown. Scale bars 10 µM. (**E, F**) LDH release (E) and NINJ1 oligomerization (Native-PAGE) (F) from WT iPSDMs with or without 5 µM Cytochalasin D pretreatment and infected with Mtb mc²6206 or Mtb mc²6230, MOI 20 for 4h. Data are means (bars) ± s.e.m. from N = 3-7 independent experiments, each with three replicates per condition. *P<0.05; **P<0.01; ****P<0.0001 by RM two-way Anova with Holm-Šídák’s multiple comparisons test against the infected sample.

### NINJ1-mediated PMR is independent of calcium and cell swelling in Mtb-infected macrophages

PM damage or pore formation can trigger ion and water fluxes across the membrane, and recent studies have reported conflicting roles for calcium fluxes and osmotic swelling in the induction of NINJ1 oligomerization and PMR(40, 45, 49). Borges et al.(49) report that Ca²⁺ fluxes are both necessary and sufficient for NINJ1-dependent PMR during pyroptosis and ATP-induced cytolysis, acting through phospholipid scrambling. To assess calcium’s role in Mtb-induced cell lysis, iPSDMs were infected with Mtb mc²6206 in calcium-free medium with the addition of BAPTA-AM (intracellular calcium chelator) or EGTA (extracellular calcium chelator). Calcium depletion did not inhibit NINJ1 oligomerization or PMR in Mtb-infected iPSDMs and rather enhanced LDH-release (Fig. 5A, B). The opposite was true for pyroptosis where calcium-depletion, specifically EGTA treatment, reduced LDH-release and slightly lowered NINJ1 oligomerization (Fig. 5C, D). Calcium chelation did not significantly reduce LDH-release from ferroptotic cells (Fig. S8A), further underscoring pathway-specific differences in NINJ1 activation.

**Figure 5:**
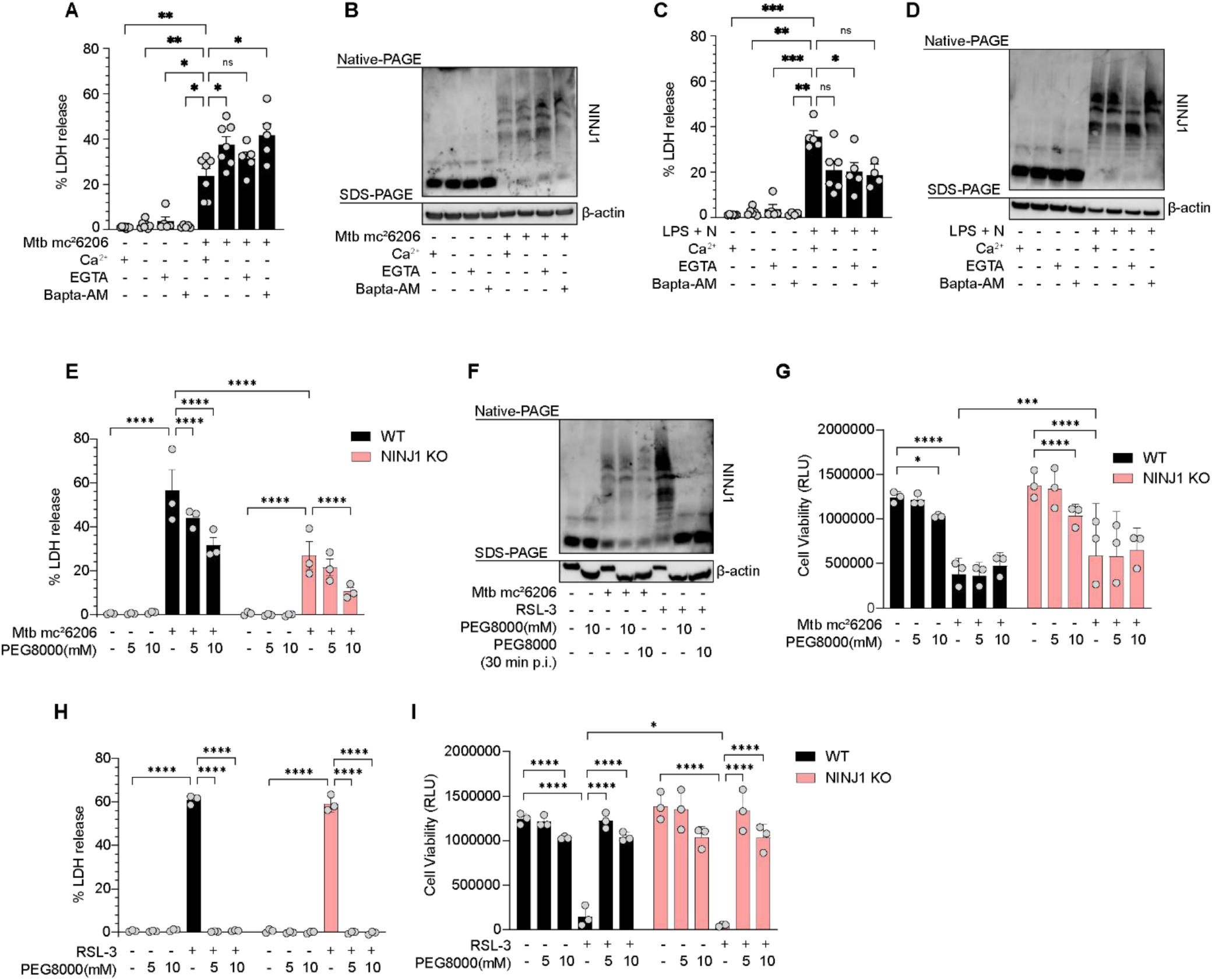
Mechanism of NINJ1 activation by Mtb. (**A-D**) LDH release (A, C) and NINJ1 oligomerization (native-PAGE) (B, D) from WT iPSDMs in DMEM with or without calcium or treated with EGTA (4 mM) or BAPTA-AM (5 µM) both in calcium-free DMEM for 30 min before infection with Mtb mc²6206, MOI 20 for 4h (A, B) or stimulation with LPS + Nigericin (C, D). Data are means (bars) ± s.e.m. of N = 5-7 independent experiments (circles), each with three replicates per condition. *P<0.05; **P<0.01; ****P<0.0001 by one-way Anova with Dunnett’s multiple comparisons test against the infected/stimulated sample with calcium. (**E-I**) LDH release (E, H) and cell viability (G, I) from WT and NINJ1 KO iPSDMs treated with PEG8000 (5 mM or 10 mM) for 30 min before infection with Mtb mc²6206, MOI 20 (E, G) or stimulation with RSL-3 (H, I) for 4h. Data are means (bars) ± s.e.m. of N = 3 independent experiments (circles), each with three replicates per condition. *P<0.05; **P<0.01; ****P<0.0001 by RM two-way Anova with Tukey’s multiple comparisons test of treatments within each cell line and stimuli between cell lines. (F) NINJ1 oligomerization (native-PAGE) from WT iPSDMs treated with PEG8000 (5 mM or 10 mM) for 30 min before infection with Mtb mc²6206, MOI 20 or stimulation with RSL-3 for 4h.

Cell swelling was proposed by Kayagaki et al.(38) to contribute to NINJ1-mediated PMR downstream of GSDMD pore formation, and osmoprotectants like sucrose or high-Mw PEGs have been shown to inhibit ferroptosis(67), possibly by restricting mechanical strain and blocking NINJ1 lesion opening(40, 45).We next assessed the effect of sucrose and PEG8000, on NINJ1-mediated PMR upon cell death and Mtb-infection in iPSDMs. Sucrose (300 mM) blocked LDH-release during ferroptosis but had no effect on Mtb-infected iPSDMs, however, sucrose itself also led to spontaneous lysis of the iPSDMs (Fig. S8B). PEG8000 reduced LDH release from Mtb-infected WT iPSDMs without affecting NINJ1 oligomerization or bacterial uptake (Fig. 5E, F and Fig. S8C, D). However, despite the expected reduction in LDH release in NINJ1 KO cells (∼50% of WT), PEG8000 reduced LDH release to a similar relative extent in both NINJ1 KO and WT iPSDMs (Fig. 5E). Cell viability was significantly and similarly reduced in WT and NINJ1 KO iPSDMs during Mtb infection and not further affected by PEG8000 (Fig. 5G). The highest concentration of PEG8000 tested (10 mM) slightly reduced the viability of untreated/uninfected iPSDMs (Fig. 5G). These data indicate that PEG8000 inhibits a residual, NINJ1-independent lytic pathway during Mtb infection and that cell swelling does not contribute to NINJ1-dependent PMR. In contrast, PEG8000 abolished LDH release and NINJ1 oligomerization in RSL-3-treated WT and NINJ1 KO iPSDMs (Fig. 5F, H and Fig. S8E, F). Surprisingly, PEG8000 also fully restored the viability of both genotypes (Fig. 5I). Pyroptosis induction with LPS+Nigericin in WT iPSDMs showed a similar pattern to Mtb infection, with PEG8000 reducing LDH release without affecting NINJ1 oligomerization; however, this was not assessed in NINJ1 KO iPSDMs (Fig. S8F, G).

## DISCUSSION

Mtb-infection drives (and inhibits) multiple modes of regulated cell death in macrophages(13, 14), with lytic death proposed to promote bacterial replication and dissemination(13, 30, 65, 68). Here, we identify NINJ1 as an executor of PMR in Mtb-infected macrophages. NINJ1 oligomerization and PMR requires the Mtb ESX-1 secretion system and occurs independently of phagocytosis or the canonical RCD pathways apoptosis, pyroptosis, necroptosis, and ferroptosis, suggesting that NINJ1 activation results from PM perturbations inflicted by ESX-1 dependent virulence factors. Cytokine release was broadly unaffected by NINJ1, except for CXCL10, indicating that NINJ1’s primary role is in cell lysis which contributes to inflammation by DAMP release and can promote Mtb dissemination. This makes NINJ1 an attractive target for adjunct host-directed therapies in tuberculosis.

Our data suggest that plasma membrane perturbation/damage serves as the initial trigger for NINJ1 activation - either caused directly, like for Mtb(9, 12) or bacterial pore-forming toxins(38, 54), or indirectly, via the induction of RCD pathways(38, 41). Although the precise mechanisms by which Mtb induces host membrane damage remain unresolved, several components of the ESX-1 secretion system including EsxA/B and EspB, as well as the cell-wall lipid PDIM, have been implicated(3, 4, 6–12). EsxA has long been associated with the lysis of artificial and host membranes(7, 69–72). It is secreted as a heterodimer with EsxB, in which EsxB is thought to mediate surface binding, while EsxA interacts directly with host membranes(69, 70, 73, 74). PDIM has been shown to facilitate the membranolytic activity of EsxA, possibly by inserting into and modifying host membranes(6–8, 11, 75). However, the mechanism underlying membrane perturbation by EsxA remains unresolved and has even been questioned(76). Other ESX-1-secreted factors may also contribute to membrane damage. EspA has been shown to regulate the secretion of EsxA and EsxB(77, 78) and while EspB can mediate cell death when the secretion of EsxA and EsxB is blocked, purified EspB did not induce macrophage cell death(12, 77, 79).

The molecular events linking PM perturbations to NINJ1 oligomerization remain undefined but likely involve biophysical changes at the membrane, such as altered lipid organization or tension. PM damage causes rapid ion-fluxes (Ca^2+^ influx, K^+^ efflux), and Ca^2+^-driven lipid scrambling was recently proposed as a trigger for NINJ1 oligomerization and PMR during pyroptosis, pneumolysin- and ATP-induced cytolysis(38, 41). A recent pre-print further suggests that caspase-induced phosphatidylserine exposure alone may be sufficient to trigger NINJ1 activation(50). Consistent with these, we observed partial inhibition of NINJ1 oligomerization and LDH-release upon Ca^2+^-chelation during pyroptosis. In contrast, NINJ1 oligomerization in Mtb-infected iPSDMs was unaffected, and Ca^2+^-chelation increased rather than decreased LDH-release, indicating that Ca^2+^-influx is not universally required for NINJ1 activation. Ca^2+^ entry through GSDMD pores, pore-forming toxins, or at contact-sites between Mtb and host membranes can also activate membrane repair mechanisms(49), and Ca^2+^-dependent membrane repair has been shown to protect cells from necrosis during Mtb-infection(45). Thus, NINJ1 activation may depend on a balance between PM damage and repair capacity. In line with this, extracellular Mtb aggregates may accumulate sufficient concentrations of virulence factors (e.g., EsxA/B) (9, 80–82) to cause local PM damage or changes in the lipid environment that trigger NINJ1 activation, whereas single or dispersed bacteria likely cause minor perturbations that are efficiently repaired. Intracellularly, similar principles may apply when phagosomal damage exceeds repair capacity(9, 35). However, it remains unclear whether NINJ1 activation in this context is triggered by PM perturbations caused by contact with cytosolic Mtb or secreted virulence factors, or indirectly through a host pathway that operates independently of caspases, RIPK3, NLRP3, or lipid peroxidation. Mtb has been reported to secrete tuberculosis necrotizing toxin (TNT) into the cytosol of infected THP-1 cells where it induces necroptotic cell death via NAD⁺ depletion(20), but the RIPK3 independence of NINJ1 activation in our system argues against a major contribution of TNT-driven necroptosis. Mtb infection has also been linked to alternative forms of programmed necrosis. For example, excess TNF can drive programmed necrosis in zebrafish(83), and type I interferon-dependent cell death has been described in mouse macrophages(84), although inhibition of type I interferon signaling did not alter Mtb-induced death in human MDMs(85). Interestingly, a recent preprint reports findings similar to ours, that Mtb-induced necrosis in human macrophages occurs independently of apoptosis, pyroptosis, necroptosis, and ferroptosis(86). Instead, necrosis was associated with DNA release and proposed to resemble a NETosis-like pathway, albeit independent of NINJ1. Thus, exactly how Mtb-induced membrane damage is coupled to NINJ1 oligomerization remains to be determined both in low- and high-burden settings.

A compromised PM leads to water influx and osmotic swelling. A model linking NINJ1 activation to cell swelling was first proposed by Kayagaki et al.(38) and subsequently supported by multiple studies, suggesting that membrane strain can promote NINJ1-mediated PMR(40, 45, 47, 48). More recently, swelling-induced membrane tension was suggested to facilitate the opening of NINJ1 lesions during ferroptosis, while oligomerization requires a distinct upstream stimulus(45). Different from Hartenian et al.(45), we find that PMR during ferroptosis in iPSDMs is NINJ1-independent. Nevertheless, high-Mw PEG inhibited both NINJ1 oligomerization and LDH release and fully restored cell viability, suggesting that osmotic protection broadly suppresses membrane damage during these conditions. Our data further indicate that neither NINJ1 oligomerization nor lesion opening in Mtb-infected iPSDMs depends on cell swelling. Instead, treatment with high–MW PEG reduced LDH release to a similar relative extent in both wild-type and NINJ1-deficient cells, implicating cell swelling as a parallel, NINJ1-independent contributor to membrane rupture in Mtb-infected macrophages. Accordingly, the contribution of NINJ1 to cell death appears highly context dependent, with different death stimuli and cell types exhibiting variable reliance on NINJ1-mediated PMR(37, 41, 87). Consistent with this, we found that NINJ1 oligomerized across all examined cell death pathways in human macrophages. Nevertheless, the lack of requirement for NINJ1 in PMR during RSL3-induced ferroptosis and zBT-induced necroptosis likely reflects the existence of alternative PMR executors, such as SIGLEC12 for necroptosis(39). In addition, NINJ1 expression levels vary between cell types and can influence membrane susceptibility to rupture(54). However, iPSDMs expressing endogenous NINJ1 were more susceptible to Mtb-induced death than THP-1 and iBMDM NINJ1 KO lines overexpressing human NINJ1-mNG, despite comparable responses to LPS+Nigericin. This suggests that membrane properties beyond NINJ1 abundance may modulate NINJ1 activation and PMR.

Mtb activates pattern recognition receptors and RCD pathways that contribute to shaping the overall inflammation and infection outcome. For example, IL-1β-release was unaffected by NINJ1 inhibition but abolished by MCC950, confirming our previous findings that NLRP3-dependent pyroptosis is induced in a fraction of Mtb-infected macrophages and solely responsible for IL-1β-production(9). Overall, deletion or inhibition of NINJ1 did not significantly affect cytokine release from Mtb-infected human macrophages except CXCL10. Like most cytokines, CXCL-10 is synthesized and trafficked through the ER-Golgi pathway and released via exocytosis(88), but our data raises the question whether there are non-canonical secretion routes for CXCL10 that involve NINJ1. NINJ1 may, however, confer a selective advantage to Mtb by facilitating bacterial escape and spread. Mtb has been shown to exploit the nutrient-rich environment of necrotic cells to replicate, form aggregates or cords, and subsequently spread to neighboring cells(30, 65, 68). Consistent with this, the extent of host cell lysis scales with bacterial burden and aggregation state(9, 12, 63, 65, 89). We find that both low- and high-burden infection induce NINJ1-dependent PMR, albeit with different kinetics, supporting a role for NINJ1 as a key mediator of inflammatory cell lysis that may facilitate bacterial dissemination.

## MATERIALS AND METHODS

### Cell culture

THP-1 monocytes were cultivated in RPMI 1640 (Sigma-Aldrich, R8758) supplemented with _L_-glutamine (Sigma-Aldrich, G7513), 10 mM HEPES (Gibco, 15630-056), sodium pyruvate (Sigma-Aldrich, S8636), D-glucose (Sigma-Aldrich, G8769) and 10% fetal bovine serum (FBS) (Gibco, A5256701). THP-1 monocytes were differentiated to macrophages using 10-50 ng/ml PMA (Sigma-Aldrich, P15859) for 1-3 days followed by one day rest before experiments. THP-1 NINJ1 KO cells were bought from Cyagen (CA, USA). iBMDMs were cultured in DMEM (EuroClone, ECM0728) with _L_-glutamine, 10% FBS and 1% Penicillin-Streptomycin (P/S) (Sigma-Aldrich, P0781), and detached using 2,5 mM EDTA (Biochrom, L2113) in PBS. iBMDM NINJ1 KO cells were generated previously(90); clonal cell lines were generated by single-cell cloning. Induced pluripotent stem cell-derived monocytes were obtained and harvested as described previously(54). Monocytes were harvested from embryonic bodies, seeded in tissue culture plates and differentiated into macrophages (iPSDMs) in RPMI 1640 supplemented with _L_-glutamine, 10% FBS and 100 ng/ml M-CSF (Peprotech, 300-25) for 5-7 days. For microscopy experiments, cells were seeded in glass-bottom 96-well plates (Cellvis, P96-1.5H-N).

### Generation of NINJ1 and GSDMD KO iPSDMs

NINJ1 and GSDMD KO iPSC cell lines were generated using CRISPR/Cas9 technology. For each gene, two single guide RNAs (sgRNAs) were designed and selected using the online design tool CRISPOR (http://crispor.tefor.net/). Nucleofection of iPSCs was performed using the P3 primary cell 4D-Nucelofector® X kit (Lonza, V4XP-3024) and program CA167. For each nucleofection reaction 2 µL (100 µM) of each synthetically chemically modified sgRNA (Synthego) (Table S1) were combined with 4 µL of S.p. Cas9 Nuclease V3 (IDT, 1081058). Following nucleofection, single cell-derived colonies were manually picked and expanded. Colonies were screened using PCR-amplification and gel electrophoresis with primers spanning the editing site (Table S2). Gene edited clones were verified by Sanger sequencing and western blot. Following editing, iPSCs were differentiated to iPSDMs.

### Lentiviral transduction of NINJ1 KO THP-1 and iBMDM with human NINJ1-mNG

Lentiviral expression vectors were made with Gateway cloning. Sequences of human NINJ1 flanked with attB sites (GeneArt gene synthesis, ThermoFisher) were cloned into pEntry vectors, which were thereafter verified by Sanger sequencing (Eurofins). A pEntry vector containing L5-L2-flanked mNeonGreen and a pLex307 destination vector with blasticidin resistance was produced in previous work (9). Expression vectors for NINJ1-mNG as were made through two-fragment Gateway recombination between pEntry-L1-NINJ1-R5, pEntry-L5-mNeonGreen-L2 and pLex307-Blast. Lentiviruses containing the NINJ1 expression vectors were produced in HEK293T cells using Lenti-X Packaging Single Shots (Takara Bio, 631275) and concentrated with Lenti-X Concentrator (Takara Bio, 631232) according to the manufacturer’s instructions. Virus-containing supernatant was filtered through a 0,45 μm PES membrane filter before virus concentration was performed. THP-1 NINJ1 KO monocytes (Cyagen, CA, USA) were transduced by resuspending approximately 750 000 cells in 1 ml filtered and concentrated virus supernatant, adding 8 μg/ml polybrene, and thereafter spinoculating at 1200*g* and 32 °C for 90 minutes. Following this, the cells were resuspended in the same virus supernatant and incubated overnight. The following day, the cells were resuspended in culture medium and expanded. Transduced THP-1 monocytes were either subjected to antibiotic selection (5 μg/ml blasticidin) or fluorescence-activated cell sorting after at least one week of expansion. For transduction of the iBMDM NINJ1 KOs, the protocol was identical, but spinoculation was done in a 12-well plate due to the cells being adherent.

### Mtb strains and infection

Mtb H37Rv (ATCC #25618) was cultivated in Middlebrook 7H9 (Sigma-Aldrich, M0178) supplemented with 10% Middlebrook OADC (BD Difco, 211886), 0.2% glycerol (Sigma Aldrich, G5516), 0.05% Tween-80 (Sigma Aldrich, P4780) or 0.05% Tyloxapol (Sigma Aldrich, T8761). 0.1 mM Sodium propionate (Sigma-Aldrich, 18108) was added to sustain PDIM levels(91). The double auxotrophic Mtb H37Rv mc^2^6206 strain (H37Rv ΔpanCD ΔleuCD)(53) and RD1-deletion mutant Mtb H37Rv mc^2^6230 strain (H37Rv ΔpanCD ΔRD1)(92) were provided by William R. Jacobs Jr. and grown like Mtb H37Rv with addition of 50 µg/mL L-leucine (Sigma Aldrich, L-8912-25G) and 24 µg/mL D-pantothenate (Sigma Aldrich, P5155-100G). Mtb mc^2^6206 was transformed to express firefly luciferase under the control of a synthetic promoter (MOP) using the pMH109 plasmid (kindly provided by David Sherman, Seattle, USA). Mtb mc^2^6206 and mc^2^6230 expressing BFP were generated as described previously(9).

For infections, log-phase Mtb cultures (OD_600_ 0.4-0.6) were centrifuged at 4754*g* for 5 min and the pelleted bacteria opsonized for 10 minutes in human serum (Blood bank of St. Olav’s Hospital, Trondheim, Norway) before dilution in cell culture medium or PBS. The resuspension was sonicated (1 min, power setting 9, VWR USC 1200 THD) and clumps of bacteria were removed by centrifugation at 300*g*, 4 min. Cells were infected with Mtb at MOI 2-30 for 4-24 hours as indicated, assuming that an OD_600_ of 1 is 3×10^6^ bacteria/ml. Intracellular bacterial loads were assessed by fluorescence microscopy (see later) or luciferase activity: cells were lysed with 5X passive lysis buffer (Promega, E1941). 50 µl lysate was mixed with 50 µl of luciferase substrate and incubated 5 min at RT before measuring luminescence using an Infinite® M plex multimode plate reader (Tecan Group Ltd., Switzerland). For prevention of phagocytosis, cells were treated with 5 µM Cytochalasin D (Tocris, 1233) before infection with Mtb mc^2^6206.

### Ligand inducers and inhibitors of cell death pathways

In iPSDMs, pyroptosis was induced by priming with 200 ng/ml LPS (*E. coli* serotype 0111:B4, Invivogen, tlrl-3pelps) for 3 hours followed by addition of 20 μM nigericin (Invivogen, tlrl-nig) for 1 hour. Apoptosis was induced with the BCL2-inhibitor, Venetoclax (125 μM, TOCRIS, 6960/10) for 4 hours. Necroptosis was induced by pretreating cells with the pan-caspase inhibitor, zVAD-FMK (25 μM, Invivogen, tlrl-vad) and the IAP-antagonist, BV-6 (10 μM, Invivogen, inh-bv6) for 30 minutes, followed by recombinant human TNF-α (10 ng/ml, PeproTech, 300–01A) for 4 hours. Ferroptosis was induced with the GPX4-inhibitor, RSL-3 (400 nM, Sigma-Aldrich, SML2234-5MG) for 4 hours. In THP-1 cells, pyroptosis was induced by treating the cells with 10 ng/ml LPS for 2-3 hours, followed by 20 µM nigericin for 1 hour. Apoptosis and ferroptosis were induced by 100 μM Venetoclax or 4 µM RSL-3 for 4 hours, respectively. In iBMDMs, pyroptosis was induced by treating cells with 0,5-1 μg/ml LPS for 2-3 hours, followed by 20 µM nigericin for 1-2 hours. Ferroptosis was induced by 4 µM RSL-3. When relevant, DRAQ7 (Biolegend, 424001) was added at a concentration of 150 nM before imaging.

Cells were pretreated with RCD inhibitors for at least 15 minutes before stimuli (ligands or Mtb) were added: NLRP3-dependent pyroptosis by 10 µM MCC950 (Sigma, PZ0280); necroptosis by 5 µM of the RIPK3 inhibitor, GSK872 (BioVision, 2673); ferroptosis by 100 nM Ferrostatin-1 (Sigma-Aldrich, SML0583-5MG). Glycine (Millipore, 104201) was used at a concentration of 33,3 mM for inhibition of NINJ1 oligomerization. For inhibition of cell swelling, cells were pretreated with 5-10 mM PEG8000 (VWR Chemicals, 0159) or 300 mM sucrose (VWR Chemicals, 27480.294) for 30 min before addition of stimuli. For inhibition of calcium-fluxes, cells grown in DMEM with calcium (Thermo Fischer Scientific, 11960044) or DMEM without calcium (Thermo Fischer Scientific, 21068028) were pretreated for 30 min with 4 mM EGTA (Sigma-Aldrich, 03777) or 5 µM BAPTA-AM (Tocris, 2787). For inhibition of palmitoylation, cells were pretreated for at least 30 min with 20 µM 2-Bromohexadecanoic acid (2-BP, Sigma-Aldrich, 21604).

### LDH assay

Cell lysis was assayed by measuring LDH-release using the CyQUANT™ LDH Cytotoxicity Assay (Invitrogen, C20301) or the LDH Cytotoxicity Detection Kit (Takara Bio, MK401), both according to the manufacturer’s instructions. In all experiments where LDH was assessed, serum levels were reduced to 1-5%. Cytotoxicity is expressed as the percentage of total LDH-release by normalizing LDH in supernatants to untreated controls and to total LDH-release in cell lysates for the various cell lines. For PEG experiments, cytotoxicity is expressed as percentage of total LDH-release normalized to untreated controls and to the total LDH-release in the cell lysates for the various PEG concentrations in the various cell lines.

### Cell viability assay

The cell viability was measured using the CellTiter-Glo® Luminescent Cell Viability Assay (Promega, G7572) according to the manufacturer’s protocol. Briefly, cells were plated in 96-well plates and treated with inhibitors and stimuli (ligands or Mtb). At the end of the experiment 100 µL CellTiter-Glo® Reagent was added to the wells. Plate was shaken on an orbital shaker for 2 min protected from light. Luminescence signal was stabilised for 10 min at RT and protected from light before measuring luminescence using an Infinite® M plex multimode plate reader (Tecan Group Ltd., Switzerland).

### Native-PAGE, SDS-PAGE and western blot

Native-PAGE, SDS-PAGE and western blot were performed as described previously(54). For Native PAGE cells were lysed using the NativePAGE Sample Prep Kit (Invitrogen, BN2008) according to the manufacturer’s instructions. A monoclonal human NINJ1 antibody (R&D Systems, MAB5105) was used to detect NINJ1. For SDS-PAGE cells were lysed with RIPA buffer (50 mM Tris-HCl pH 7, 150 mM NaCl, 1% Triton X-100, 0,1% SDS) or native page lysates were used. NINJ1 was detected by Western blotting using a polyclonal human NINJ1 antibody (R&D Systems, AF5105). Anti-β-actin (Cell Signaling Technology, 8457 or 12620) was used for all loading controls.

### ELISA and Multiplex Cytokine analysis

IL-1β was measured in the supernatant using Human IL-1beta/IL-1F2 DuoSet ELISA (R&D Systems, DY201) according to the manufacturer’s instructions. Multiplex ELISA was performed using custom-made Human ProcartaPlex Mix&Match panels of 17, 20 or 23 cytokines/chemokines according to the manufacturer’s instructions (Invitrogen).

### Fluorescence microscopy

All imaging was performed with a Zeiss TIRF III (Carl Zeiss GmbH), using an alpha Plan-Apochromat 63×/1.46 oil-immersion objective, occasionally employing a 1,6× tube lens. A 37 ℃ and 5% CO_2_ incubation chamber was utilized in all live-cell imaging. Furthermore, all images were captured with an ORCA-Fusion CMOS camera (Hamamatsu, C14440-20UP). For live-cell imaging, cells were cultured in a 96-well plate with glass bottom (Cellvis, P96-1.5H-N) over-night (iBMDMs) or for 3 days (THP-1s) in culture medium. Right before the experiment, cells were washed 1x with culture medium and DRAQ7 (ThermoFisher, D15106) was added to monitor cell permeabilization. Afterwards, the cells rested at the microscope at 37 ℃ and 5% CO_2_ for 30 min before saving 2-3 positions per well that show several cells. Images of cells were taken before treatment with inhibitors, RCD ligands or infection and the same cells were imaged throughout the whole experiment. Cell morphology was observed with bright field, cell permeabilization was monitored by DRAQ7 uptake upon illumination with a 647 nm laser with epi fluorescence, NINJ1-NeonGreen dynamics were monitored by excitation with a 488 nm laser in TIRF and Mtb bacteria were illuminated with a 405 nm laser in widefield.

### Image analysis

Image processing and quantification were done in FIJI(93). To quantify the number of NINJ1-mNG puncta, either TrackMate v7.11.1 or the “Find maxima” tool with manually adjusted prominence values were used on a single-cell basis. For kinetics measurements, the quantifications were thereafter aligned manually to the frame where NINJ1-mNG clustering started and normalized to their maximum count in each cell. The increase of NINJ1 puncta was fitted with a 4P sigmoidal fit in GraphPad Prism versions 9 to 10.5.0 (GraphPad Software LLC). The DRAQ7 influx was quantified by using the *measure* tool on a region of interest (ROI) that fit into the nucleus for all images in the time-lapse to obtain the mean gray value of the ROI for all the images. The measurements were thereafter aligned as described above, the background was subtracted and all intensity values were normalized to the mean gray value of the first frame. To quantify the NINJ1-mNG puncta density per cell in TIRF microscopy images, ROIs were manually drawn around cells, and the *find maxima* command was used within them. Thereafter, the areas of the ROIs were determined with the *measure* tool and the puncta per squared micron was calculated for each cell. The DRAQ7 influx and bacterial load per cell were measured by using the *measure* tool on a region of interest (ROI) that fit into the nucleus (DRAQ7) or the whole cytosol (Mtb mc^2^6206 or mc^2^6230 BFP) and assess the overall intensities. All values were normalized to the results of Mtb mc^2^6206 infection at 0 min.

## Statistical analyses, graphs and Figures

All graphs were produced, and statistical analyses were performed in GraphPad Prism versions 9 to 10.5.0 (GraphPad Software LLC). Figures were compiled using Adobe Illustrator 2025 version 29.5.1 (Adobe).

## Acknowledgements

We thank Unni Nonstad and Linn-Karina Selvik for technical help in this work, Alberto Diez Sanchez at CMIC for help with image analysis, and prof. Thierry Soldati, Universite de Geneve, Switzerland for discussions/advice. All imaging was performed at the Cellular and Molecular Imaging Core Facility (CMIC), Norwegian University of Science and Technology (NTNU).

## Funding

Research Council of Norway grant 287696 (THF)

Research Council of Norway grant 223255 (THF)

The Olav Thon Foundation (THF)

Norwegian University of Science and Technology (NTNU) (THF)

European Union’s Horizon Europe and innovation programme under the Marie Skłodowska-Curie grant agreement No 101211876 (SK)

CMIC is funded by the Faculty of Medicine and Health Sciences at NTNU, Central Norway Regional Health Authority and NALMIN II (NFR 322607)

## Author contributions

THF conceptualized and supervised the project and acquired funding. RSRS, MH, SK, MB, AM, LR and KSB performed experiments and analyzed the data. PD, MH and RSRS generated cell lines. RSRS, MH and SK prepared the figures. RSRS, MH, SK and THF wrote the original manuscript draft; PD, JCK, KF and KSB reviewed and edited the manuscript.

## Data, code, and materials availability

Source data including unprocessed gels and blots, statistics source data and image quantifications used in plotted graphs will be provided with the published manuscript in an Excel Workbook. Imaging source data with metadata and image analysis pipelines will be made available in public repositories (and linked here).

