## Supplementary Figures for "NINJ1 mediates necrosis of *Mycobacterium tuberculosis* infected human macrophages"

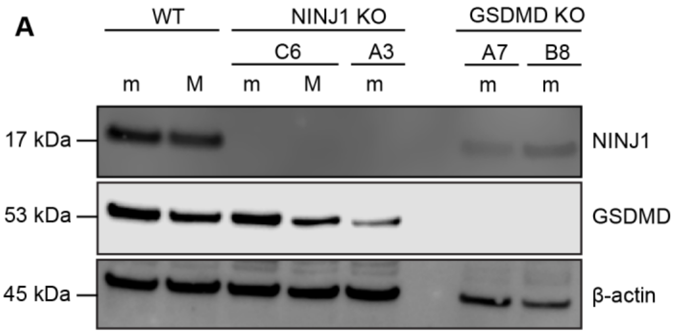


**Fig. S1. NINJ1 and GSDMD KO verification.**

(**A**) Lysates from WT, NINJ1 KO and GSDMD KO iPSC-derived monocytes (m) and macrophages (M) subjected to SDS-PAGE and immunoblotted for NINJ1, GSDMD and β-actin. A3 and C6 are NINJ1 KO clones; A7, B8 are GSDMD KO clones.

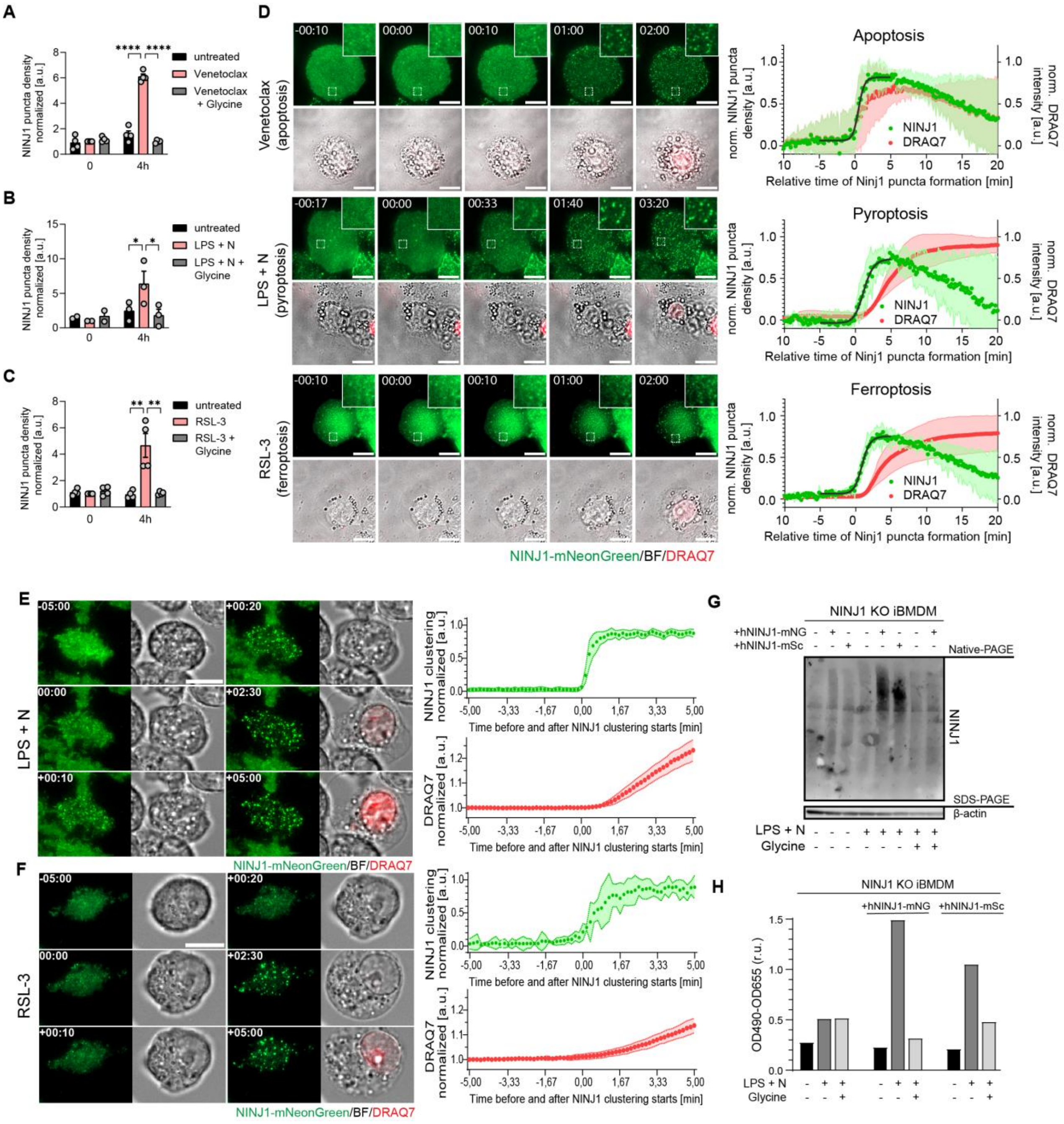


**Fig. S2. NINJ1 clustering/puncta densities and cell death kinetics during apoptosis, pyroptosis and ferroptosis.**

(**A-C**) Quantification of NINJ1-mNG puncta densities after 0 and 4 h of cell death stimuli: (A) apoptosis – Venetoclax; (B) pyroptosis – LPS + N; (C) ferroptosis – RSL-3, with or without glycine treatment in THP1-NINJ1-mNeonGreen cells. Data are means (bars) ± s.e.m from N = 3-4 independent experiments (circles), each including analysis of 4-15 cells per condition. Significant differences are indicated as **P*<0.05; ***P*<0.01; *****P*<0.0001 by RM two-way Anova with Tukey's multiple comparisons test. (**D**) Kinetics of NINJ1 oligomerization and DRAQ7 intensity in THP-1 NINJ1-mNG cells after treatment with different cell death stimuli (Venetoclax, RSL-3, LPS+N) monitored with frame rates of 8-10 sec with representative time-lapse images (Time in sec) of NINJ1 puncta formation (TIRF, green) and DRAQ7 (WF, red) aligned by start of NINJ1 clustering. Scale bars 10 µm. Intensity values were normalized to the highest value for each cell and aligned by the start of NINJ1 oligomerization. Data are means ± s.d. with sigmoidal fits for NINJ1-mNG puncta density increase. N = 3 experiments, 8-12 cells per condition. (**E, F**) Time-lapse microscopy of NINJ1 KO immortalized mouse BMDMs reconstituted with mNG-tagged human NINJ1 treated with LPS + nigericin (E) or RSL-3 (F). Image panels show NINJ1-mNG oligomerization (TIRF, green), DRAQ7 influx (red, WF) and BF with quantification of NINJ1-mNG puncta density kinetics and increase of DRAQ7 intensities aligned by start of NINJ1 clustering. Time points shown in upper left corner in min. Scale bar 10 µm. Quantification shown to the right, data are presented as mean ± 95% CI. (**G, H**) NINJ1 oligomerization (Native-PAGE) (G) and LDH release (H) from mouse NINJ1 KO iBMDMs expressing human NINJ1-mNG or NINJ1-mSc treated with LPS + nigericin with or without glycine treatment.

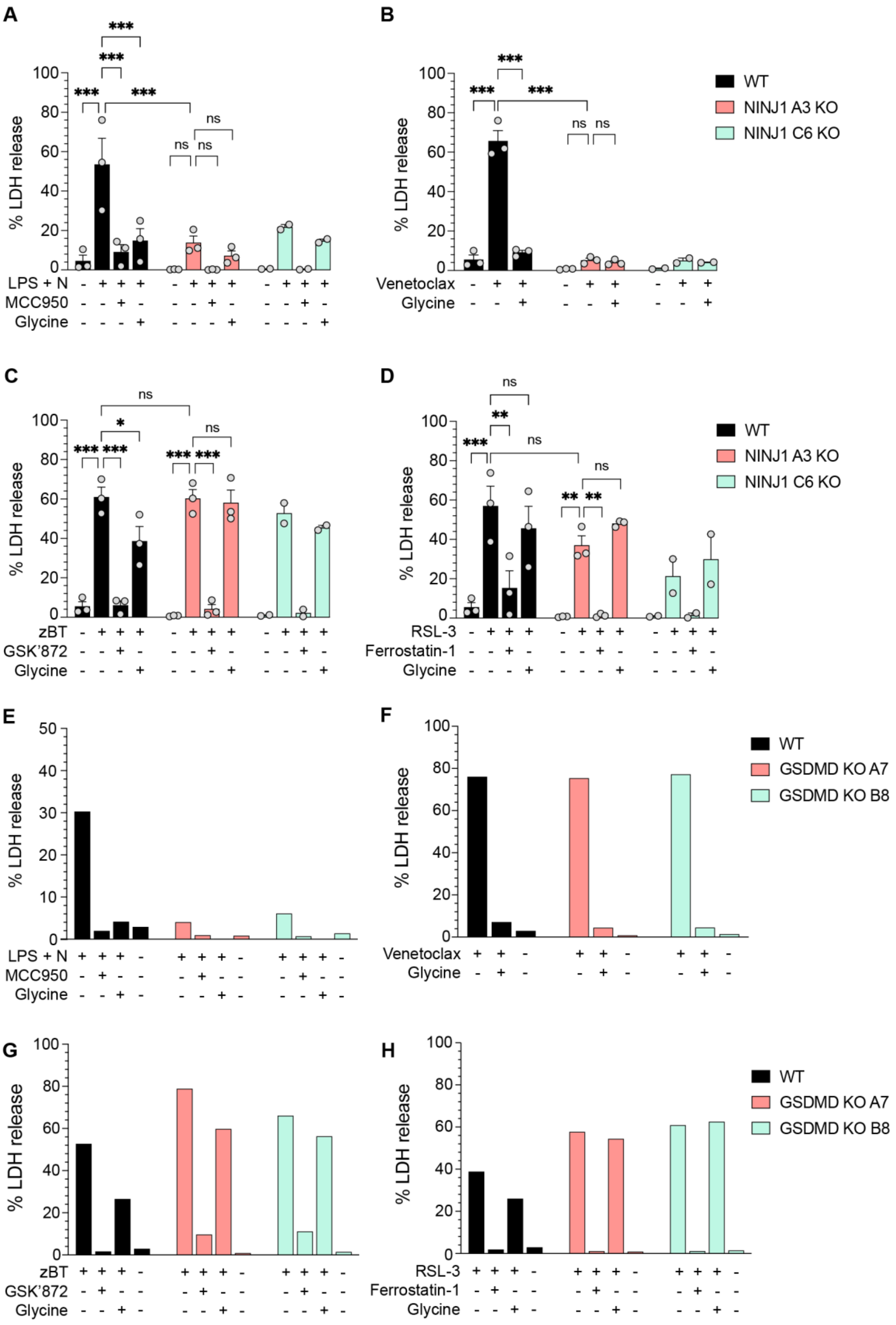
A
% LDH release
100
80
60
40
20
0
***
***
***
***
ns
ns
ns
LPS + N - + + + - + + + - + + +
MCC950 - - + - - - + - - - + -
Glycine - - - + - - - + - - - +
B
% LDH release
***
***
***
ns
ns
Venetoclax - + + - + + - + +
Glycine - - + - - + - - +
WT
NINJ1 A3 KO
NINJ1 C6 KO
C
% LDH release
ns
*
***
***
***
***
ns
zBT - + + + - + + + - + + +
GSK'872 - - + - - - + - - - + -
Glycine - - - + - - - + - - - +
D
% LDH release
ns
**
***
ns
**
**
ns
RSL-3 - + + + - + + + - + + +
Ferrostatin-1 - - + - - - + - - - + -
Glycine - - - + - - - + - - - +
WT
NINJ1 A3 KO
NINJ1 C6 KO
E
% LDH release
50
40
30
20
10
0
LPS + N + + + - + + + - + + + -
MCC950 - + - - - + - - - + - -
Glycine - - + - - - + - - - + -
F
% LDH release
Venetoclax + + - + + - + + -
Glycine - + - - + - - + -
WT
GSDMD KO A7
GSDMD KO B8
G
% LDH release
zBT + + + - + + + - + + + -
GSK'872 - + - - - + - - - + - -
Glycine - - + - - - + - - - + -
H
% LDH release
RSL-3 + + + - + + + - + + + -
Ferrostatin-1 - + - - - + - - - + - -
Glycine - - + - - - + - - - + -
WT
GSDMD KO A7
GSDMD KO B8

**Fig. S3. Plasma membrane rupture during pyroptosis, apoptosis, necroptosis and ferroptosis in WT, NINJ1 KO and GSDMD KO iPSDMs.**

(**A-H**) LDH-release from iPSDM WTs and NINJ1 KO clones A3, C6 (A-D) or GSDMD KO clones A7, B8 (E-H) pretreated with or without glycine or inhibitors as indicated and treated with or without cell death stimuli for 4h: (A, E) pyroptosis – LPS and nigericin (LPS+N) w/wo NLRP3 inhibitor MCC950; (B, F) apoptosis – venetoclax; (C, G) necroptosis – Z-VAD-FMK+BV-6+TNF-α (zBT) w/wo RIPK3 inhibitor GSK'872; (D, H) ferroptosis – RSL-3 w/wo Ferrostatin-1. (A-D). Data are means (bars) ± s.e.m. of N = 3 (A3) or N = 2 (C6) independent experiments (circles), each with three replicates per condition. *$P<0.05$; **$P<0.01$; ****$P<0.0001$ by RM two-way Anova with Tukey's multiple comparisons test within and between WT and NINJ1 A3 KO. (E-H) Data are means of two or three technical replicates from N = 1 independent experiment.

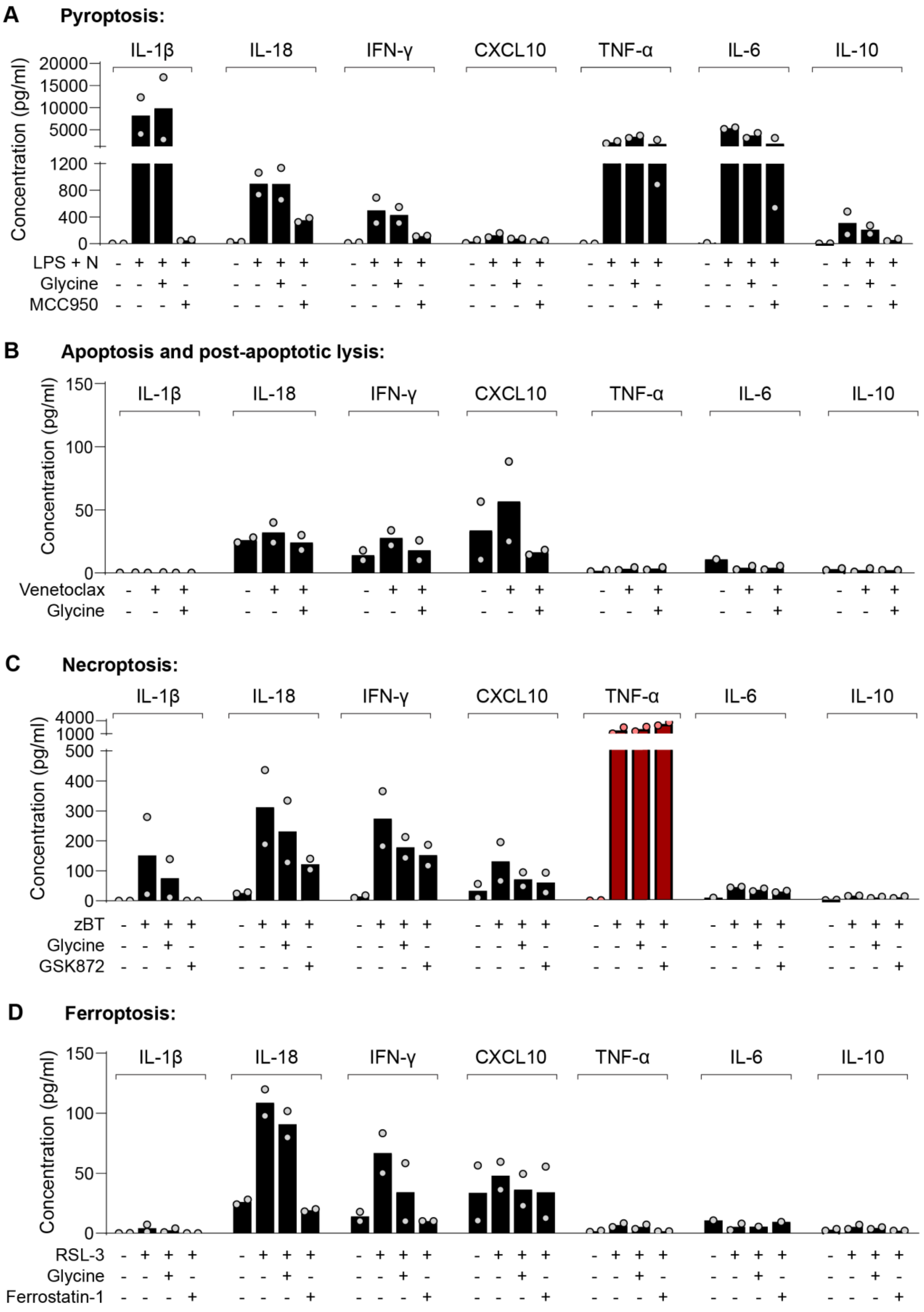
A Pyroptosis:
IL-1β
IL-18
IFN-γ
CXCL10
TNF-α
IL-6
IL-10
Concentration (pg/ml)
20000
15000
10000
5000
1200
800
400
0
LPS + N
Glycine
MCC950
B Apoptosis and post-apoptotic lysis:
150
100
50
Venetoclax
C Necroptosis:
4000
1000
500
300
200
100
zBT
GSK872
D Ferroptosis:
RSL-3
Ferrostatin-1

**Fig. S4. Cytokine release from macrophages during different cell death modalities (pyroptosis, apoptosis, necroptosis and ferroptosis).**

(**A-D**) Cytokine release from WT iPSDMs pretreated with glycine or inhibitors as indicated for 30 min and treated with cell death stimuli for 4h: (A) pyroptosis – LPS and nigericin (LPS+N) w/wo NLRP3 inhibitor MCC950; (B) apoptosis – venetoclax; (C) necroptosis – zVAD+BV-6+TNF-α (zBT) w/wo RIPK3 inhibitor GSK'872; (D) ferroptosis – RSL-3 w/wo Ferrostatin-1. Cytokine levels were assessed by multiplex ELISA. Data are means (bars) of N = 2 independent experiments (circles), each with two technical replicates per condition.

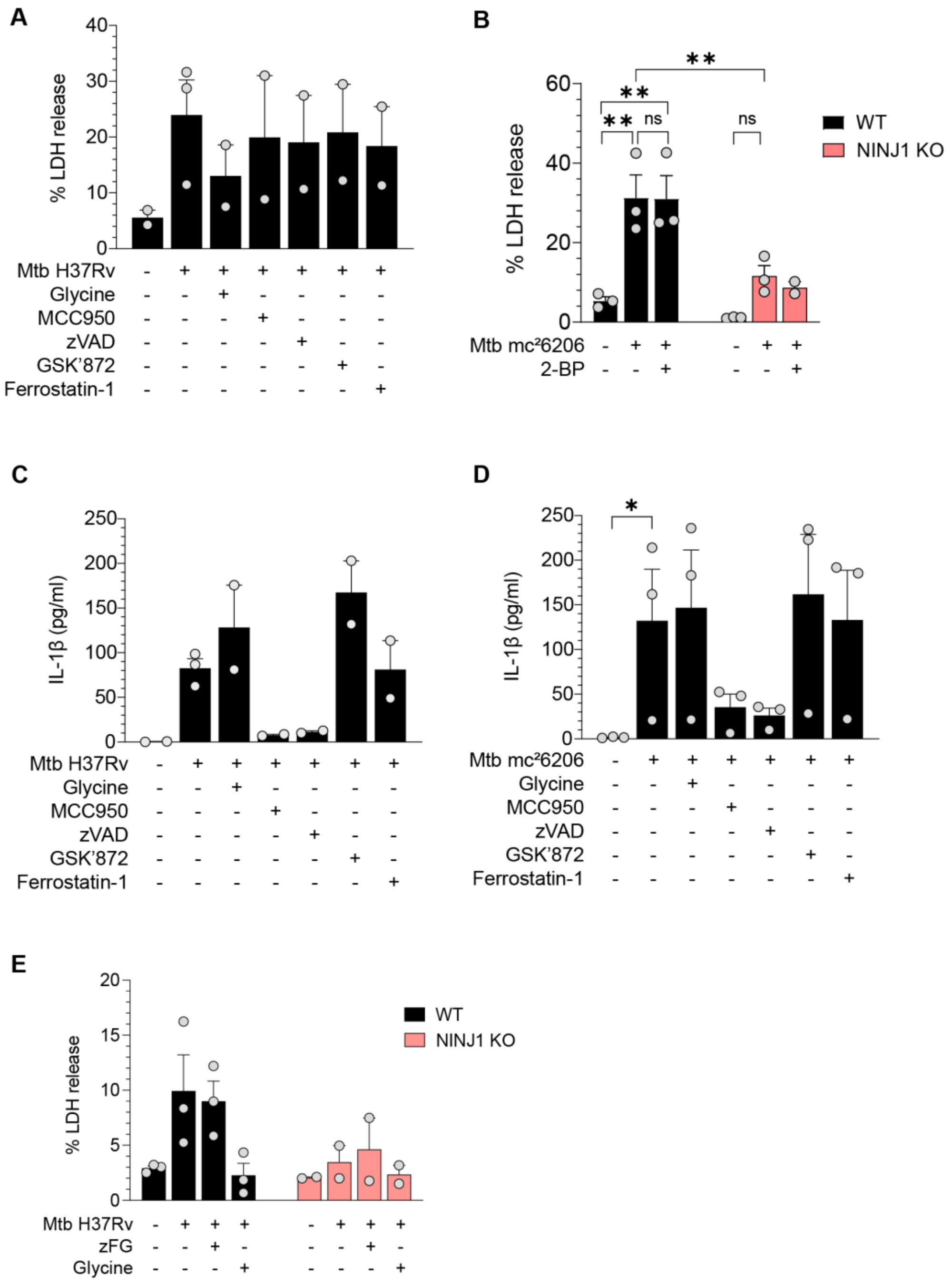


**Fig. S5. Inhibition of cell death pathways and palmitoylation does not affect plasma membrane rupture in Mtb-infected iPSDMs.**

(**A**) LDH release from WT iPSDMs pretreated with glycine and cell death inhibitors (MCC950, Z-VAD-FMK, GSK’872 and Ferrostatin-1) for 30 min and infected with Mtb H37Rv, MOI 7,5 for 24h. Data are means (bars) ± s.e.m. of N = 2-3 independent experiments (circles), each with three replicates per condition. (**B**) LDH release from WT and NINJ1 KO iPSDMs pretreated with 20 µM 2-Bromohexadecanoic acid (2-BP) for 30 min and infected with Mtb mc²6206, MOI 20 for 4h. Data are means (bars) ± s.e.m. of N = 3 independent experiments (circles), each with three replicates per condition. *$P<0.05$; **$P<0.01$; ****$P<0.0001$ by RM two-way Anova with

Šídák's multiple comparisons test of treatments within each cell line and stimuli between cell lines. All comparisons are shown. (**C, D**) IL-1β release from WT iPSDMs pretreated with glycine or cell death inhibitors (MCC950, Z-VAD-FMK, GSK'872 and Ferrostatin-1) for 30 min and infected with Mtb H37Rv, MOI 7,5 for 24h (C) or Mtb mc²6206, MOI 20 for 4h (D). Data are means (bars) ± s.e.m. of N = 2-3 independent experiments (circles), each with three replicates per condition. **P*<0.05; ***P*<0.01; *****P*<0.0001 by RM two-way Anova with Dunnett's multiple comparisons test against the infected sample (D). (**E**) LDH release from WT and NINJ1 KO iPSDMs pretreated with glycine or a cocktail of inhibitors (zFG) including Z-VAD-FMK, Ferrostatin-1 and GSK'872 and infected with Mtb H37Rv, MOI 7,5 for 24h. Data are means (bars) ± s.e.m. of N = 2-3 independent experiments (circles), each with three replicates per condition.

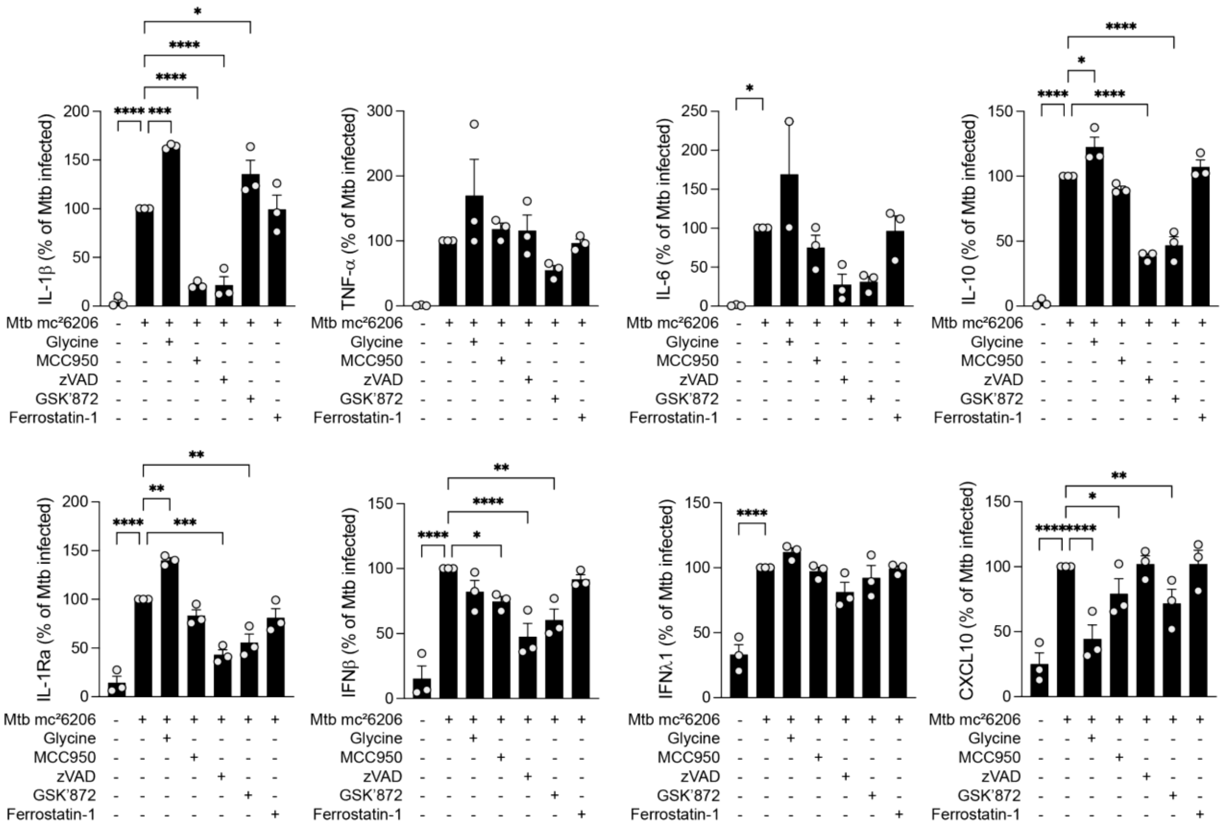


**Fig. S6. Impact of individual cell death pathways and NINJ1 on the inflammatory response induced by Mtb.**

Cytokine release from iPSDMs WT pretreated with glycine, MCC950, Z-VAD-FMK, GSK'872 and Ferrostatin-1 and infected with Mtb $mc^2 6206$, MOI 20 for 4h. Data are normalized against the Mtb $mc^2 6206$ infected sample and presented as means (bars) ± s.e.m. of N = 3 independent experiments (circles), each with three replicates per condition. **P*<0.05; ***P*<0.01; *****P*<0.0001 by RM two-way Anova with Dunnett's multiple comparisons test against the infected sample. Only significant results are shown.

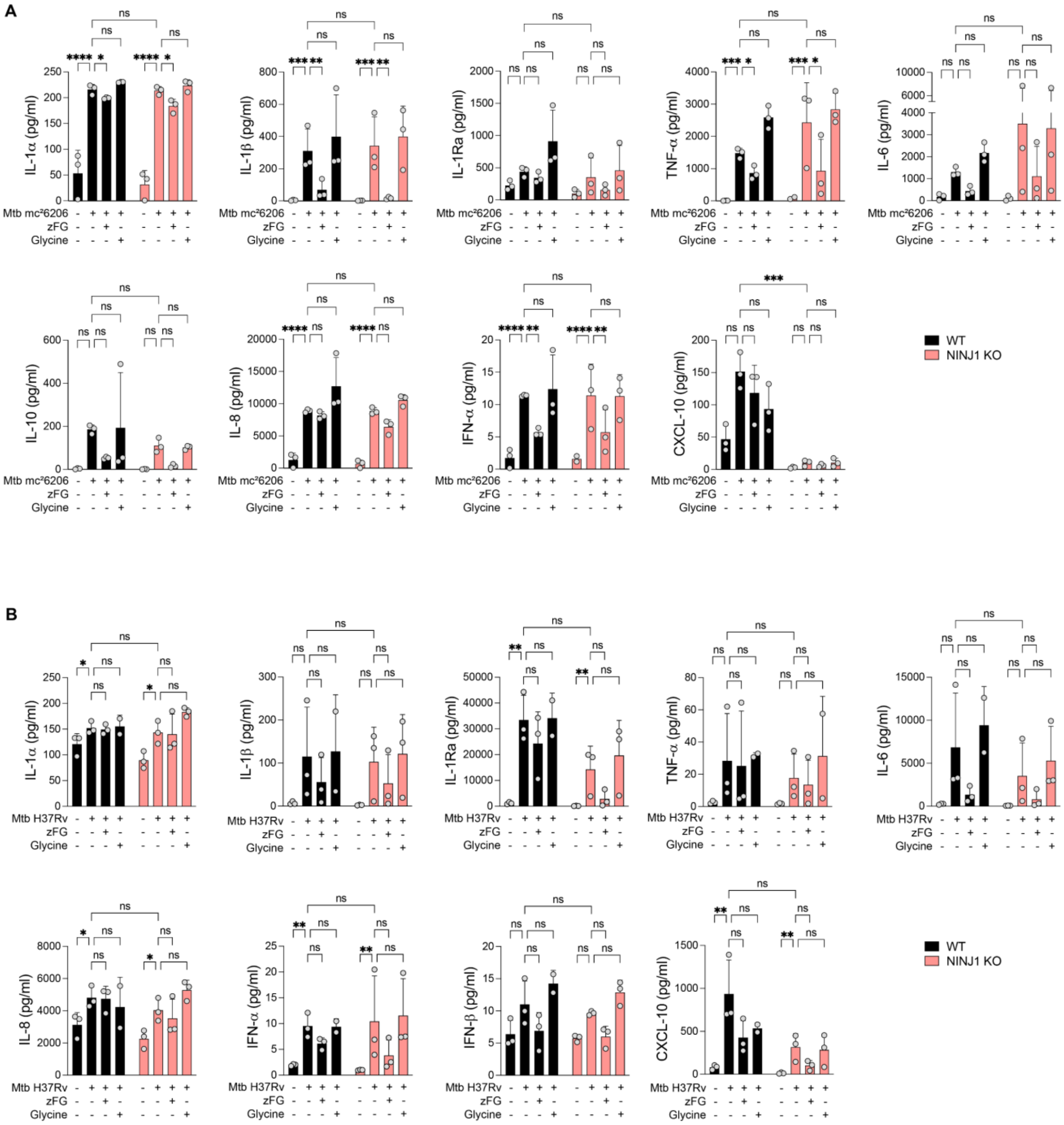


**Fig. S7. Impact of individual cell death pathways and NINJ1 on the inflammatory response induced by Mtb.**

(**A, B**) Cytokine release from WT and NINJ1 KO iPSDMs pretreated with glycine or a cocktail of inhibitors (zFG) including Z-VAD-FMK, Ferrostatin-1 and GSK'872 and infected with Mtb mc²6206, MOI 20 for 4h (A) or Mtb H37Rv, 7.5 MOI for 24h (B). Data are means (bars) ± s.e.m. of N = 3 independent experiments (circles), each with three replicates per condition. **P*<0.05; ***P*<0.01; *****P*<0.0001 by RM two-way Anova with Šídák's multiple comparisons test of treatments within each cell line and infection between cell lines. All comparisons performed are shown.

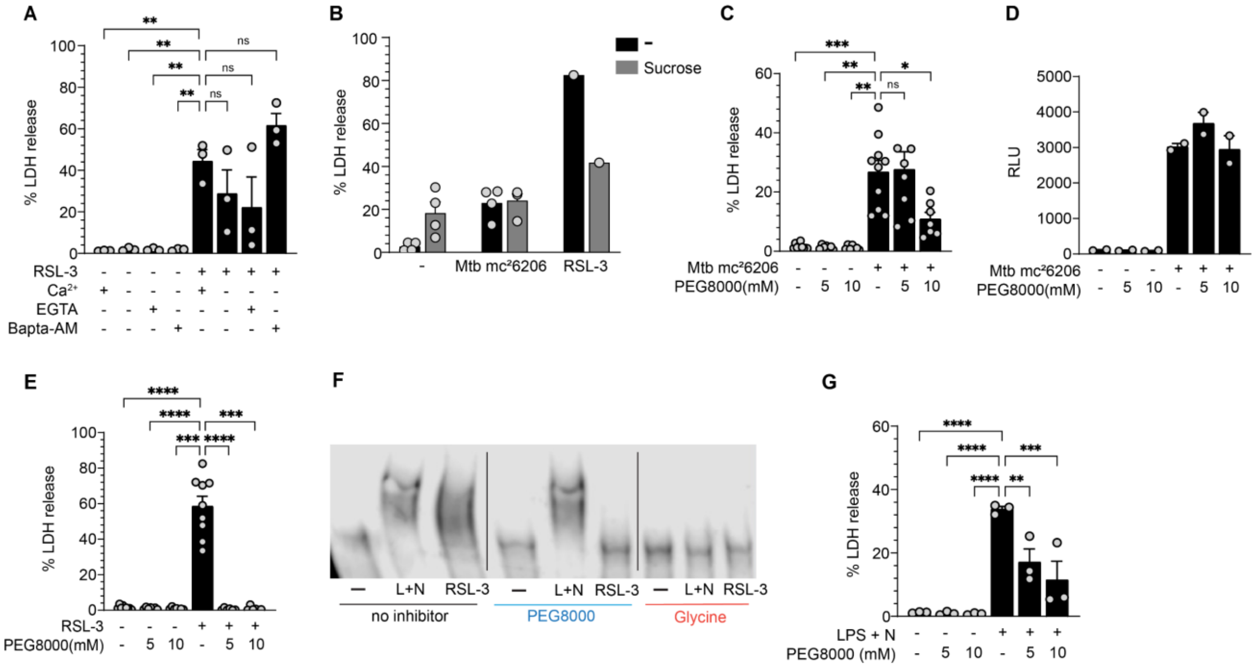


**Fig. S8. Mechanism of NINJ1 activation by Mtb.**

(**A**) LDH release from WT iPSDMs in DMEM with or without calcium or pretreated with EGTA (4 mM) or BAPTA-AM (5 µM) in calcium-free DMEM before stimulation with RSL-3 for 4h. Data are means (bars) ± s.e.m. of N = 3 independent experiments (circles), each with three replicates per condition. *$P$<0.05; **$P$<0.01; ****$P$<0.0001 by one-way Anova with Dunnett's multiple comparisons test against the stimulated sample with calcium. (**B**) LDH release from WT iPSDMs treated with 300 mM sucrose for 30 min before infection with Mtb mc²6206, MOI 20 or stimulation with RSL-3 for 4h. Data are means (bars) ± s.e.m. of N = 1-4 independent experiments (circles), each with three replicates per condition. (**C-G**) LDH release (C, E, G), Relative light units (RLU) (D) and native-PAGE (F) from WT iPSDMs treated with glycine or PEG8000 (5 mM or 10 mM) for 30 min before infection with Mtb mc²6206, MOI 20 (C, D), stimulation with RSL-3 (E, F) or LPS + Nigericin (F, G) for 4h. Data are means (bars) ± s.e.m. of N = 7-10 (C,E), N = 2 (D) or N = 3 (G) independent experiments (circles), each with three replicates per condition. *$P$<0.05; **$P$<0.01; ****$P$<0.0001 by RM one-way ANOVA with Dunnett's multiple comparisons test against the infected/stimulated sample. All comparisons performed are shown.

**Table S1.** Single guide RNA sequences used in this study.

| **Name sgRNAs** | **Sequence (5'→3')** |
|---|---|
| NINJ1 sgRNA1 | GGGCGGCCGCACCATGGACT |
| NINJ1 sgRNA2 | GAGGAGTACGAGCTCAACGG |
| GSDMD sgRNA1 | CTTGCTTTAGACGTGCAGCG |
| GSDMD sgRNA2 | TTCCACTTCTACGATGCCA |

**Table S2.** Primer pair sequences used in this study.

| **Name primers** | **Sequences (5’à3’)** |
|---|---|
| NINJ1 fw | GACGTCCCCCAACACTCTG |
| NINJ1 rev | GGGTCCTCTGGGGCCAT |
| GSDMD fw | GGCCTTTCCAAAGGTCCCTC |
| GSDMD rev | CAGGGTTGCTTGGGGTAGGT |
